# Detection of misfolded proteins in neurodegenerative disease models with a highly sensitive chemiluminescence probe

**DOI:** 10.1101/2025.04.12.648090

**Authors:** Biyue Zhu, Richard Van, Huizhe Wang, Shi Kuang, Yuntao Jia, Erick Calderon Leon, Jing Zhang, Jun Yang, Howard Hong, Fleur Lobo, Astra Yu, Johnson Wang, Rudolph E. Tanzi, Can Zhang, Xiaobo Mao, Yihan Shao, Fan Yang, Chongzhao Ran

## Abstract

Visualizing misfolded proteins would greatly facilitate early diagnosis, etiology elucidation, and therapy monitoring of neurodegeneration. Although several probes have been reported, simple and versatile detection in vivo is still challenging. We demonstrated that both generic and precise detection of misfolded proteins could be achieved with a chemiluminescence probe, ADLumin-1. For the generic aspect, ADLumin-1 was highly sensitive to various misfolded proteins, showing up to 127.73-fold higher signal-to-noise ratio than Thioflavin T. ADLumin-1 could also non-invasively visualize misfolded proteins in mouse models of Parkinson’s disease, Alzheimer’s disease and amyotrophic lateral sclerosis. Furthermore, ADLumin-1 displayed precise detection value for α-synuclein by combining with PMCA in vitro and bioorthogonal ChRET imaging technology in vivo. ADLumin-1 can selectively detect α-synuclein in CSF at the femtomolar level and enables in situ monitoring of misfolded α-synuclein in vivo. Combining generality and precision, our findings could be widely applied in preclinical and clinical studies of neurodegenerative diseases.

**Teaser:** A highly sensitive chemiluminescence probe enables both generic and precise detection of various misfolded proteins

## Introduction

In 1997, Carrell and Lomas proposed “conformational disease” as a general category for several disorders, including the most influential neurodegenerative diseases of Alzheimer’s disease (AD) and Parkinson’s disease (PD) (*1–3*). “Conformational disease” is closely associated with proteinopathy (*4*). The key feature of proteinopathy is the accumulation of proteins of abnormal conformations, such as misfolded β-sheet, that triggers downstream harmful cascades (*5*). Presently, the occurrence of neurological proteinopathies is on the rise with advancing age. Nevertheless, there remains a significant need for effective therapeutics and robust diagnostic tools (*5*).

Although comprehensive studies have been conducted, the disease etiology related to the aggregation and propagation of misfolded proteins remains elusive (*6–9*). The most widely studied peptides and proteins include β-amyloid (Aβ) (*10, 11*), tau(*12*), α-synuclein (α-syn) (*13*) amylin (*14*), polyglutamine (PolyQ) (*15*), transthyretin (TTR) (*16*), and serum amyloid A (SAA) (*17*). However, it is widely believed that the disruption of their native conformation facilitates the formation of misfolded β-sheets, and consequentially assemble into toxic oligomers, protofibrils and fibrils, and even large deposits (*7*). This aggregation process is often accompanied by propagation and has been closely associated with disease etiology. For example, Aβ and tau species could assemble into oligomers and self-propagate throughout the brain, which leads to the over-activation of microglia, neuroinflammation, and consequential synapse loss in AD (*18–20*). Dynamic monitoring of misfolded proteins could facilitate the elucidation of structural biology, and proteinopathy etiology and assist in the anti-neurodegeneration drug discovery (*5*). Thus, simple, convenient, and widely applicable detection methods are highly needed.

Notably, the Cryo-EM structures of the misfolded proteins in proteinopathies revealed that most of them share similar β-sheet structures (*21, 22*). In general, several amino acids, including Ala, Val, Ile, Leu, and Phe, can be highly frequently found in most of the β-sheets, and the tendency of these amino acids to adopt β-sheet structures was generally known from their statistical occurrence in protein secondary structures (commonly referred to as the “Chou-Fasman parameters”) (*23, 24*) and from empirical studies (*25*). Here, we termed these peptides and proteins as “misfoldon”, which share the common feature of misfolded β-sheet conformation. Although these “misfoldons” have different amino acid sequences, charge status, hydrophobicity, and water solubility, the commonality is that they could gradually form oligomers and fibrils under proper conditions in vitro and in vivo (*26, 27*). In addition, like prions, with the seed of a misfoldon, the conversion from its native conformation to misfolded β-sheet and the aggregation of resulting β-sheets can often be accelerated (*6*). From the Cryo-EM structure, we noticed one or more hydrophobic pockets that are formed by at least three amino acids from Ala, Val, Ile, Leu, and Phe (*21, 22*), suggesting that such a hydrophobic pocket is a general secondary structure for β-sheets, and it is reasonable to speculate that some generic ligands may match with this typical binding site. Indeed, benzothiazole-based fluorescence dye Thioflavin T (ThT) has been developed for monitoring the broad spectrum of the misfoldons and is widely used for in vitro diagnosis and etiology elucidation. The applications of ThT include detecting the concentrations of β-sheet fibrils in solution, biofluid, and histological slides (*28–30*). Recently, protein-misfolding cyclic amplification (PMCA) and real-time quaking-induced conversion (RT-QuIc) assay have been developed for ultrasensitive detection of prions or α-syn in biofluids and tissue samples using ThT fluorescence (*31–34*). Those methods were developed based on misfoldon propagation and have been successfully applied in the clinical diagnosis of PD and prion diseases (*35–37*). However, ThT has several stringent limitations. For example, the biological auto-fluorescence causes high background signals, and the short emission wavelength of ThT hampers deep tissue penetration restricting its applications for in vivo imaging (*28–30*).

Using advanced imaging technologies for monitoring misfoldons has raised tremendous interest (*38–40*). Numerous fluorescence imaging probes have been developed for misfoldon biomarkers, but they still face several obstacles, including high background, limited signal-to-noise ratio (SNR), and penetration depth (*41–43*). Different from fluorescence imaging, molecularly produced light, such as chemiluminescence, could significantly reduce background interference and thus facilitate deep-tissue imaging (*44–47*). However, currently none of the chemiluminescence probes have been developed for monitoring a broad spectrum of misfoldons.

Recently, we reported that ADLumin-1, a chemiluminescence probe developed by our group, could be used for imaging Aβ species in vitro and in vivo (*42*). The molecular docking result revealed that ADLumin-1 interacted with the hydrophobic pocket formed by Phe19, Ala21, Val24, and Ile31. The binding pose is similar to ThT, which binds along the long axis of Aβ fibrils. Based on the similarity of the interaction model of ADLumin-1 and ThT with misfoldons, we speculated that ADLumin-1 could be a versatile probe for various misfoldons. In this report, we revealed that ADLumin-1 could robustly respond to five or more of the most studied misfoldons and enable robust in vivo imaging of α-syn, tau, and TAR DNA-binding protein 43 (TDP-43) deposits in transgenic mice models. Despite this generality of misfoldon detection, ADLumin-1 also achieves selective misfoldon visualization by combining the PMCA technique in vitro and chemiluminescence resonance energy transfer (ChRET) in vivo. We demonstrated that ADLumin-1 could selectively detect α-syn fibrils in biofluids and monitor its accumulation during the aging of the transgenic A53T mouse model. Thus, we envisioned that ADLumin-1 could be a powerful tool for proteinopathies early diagnosis, and etiology elucidation and also assist in anti-neurodegeneration therapy.

## Results

### ADLumin-1’s specificity to β-sheet-rich structures

For the detection of protein fibrils, fluorescence probes have been routinely used in solution tests and in vivo imaging; however, chemiluminescence probes have been rarely used (*40–42*). In our previous studies, we demonstrated that a significant amplification of chemiluminescence signal could be observed with ADLumin-1 in the presence of Aβ fibrils, and the amplification is due to the turn-on effect of ADLmin-1 upon binding to the hydrophobic environment inside the β-sheet fibrils (*42*). This is further supported by molecular docking studies that suggested ADLumin-1 binds along the long axis of the β-sheet of Aβ fibrils (*42*). Interestingly and notably, the binding pose of ADLumin-1 closely resembles that of ThT when bound to Aβ fibrils (Fig. S1) (*48, 49*). This led us to hypothesize that, like ThT, ADLumin-1 might serve as a generic probe for specifically targeting β-sheet-rich structures (Fig. 1A). To investigate the specificity of ADLumin-1 for β-sheet fibrils, we used four model peptides PA-K, PA-K2, PA-E and PA-E2 (Fig. 1B and Fig. S2A) (*50*). It is well-documented that they are peptide amphiphiles (PA) and could self-assemble to form fibrils under various conditions (*50*). PA-E and PA-K, which do not have amino acids for β-sheet formation, primarily rely on electrostatic interaction to form fibrils, whereas PA-E2 and PA-K2, which contain typical amino acids (three Valines and three Alanines) for β-sheet formation, have a higher tendency to produce β-sheet fibrils via van de Waals interaction and hydrogen bonding (*50*).

**Fig. 1.**
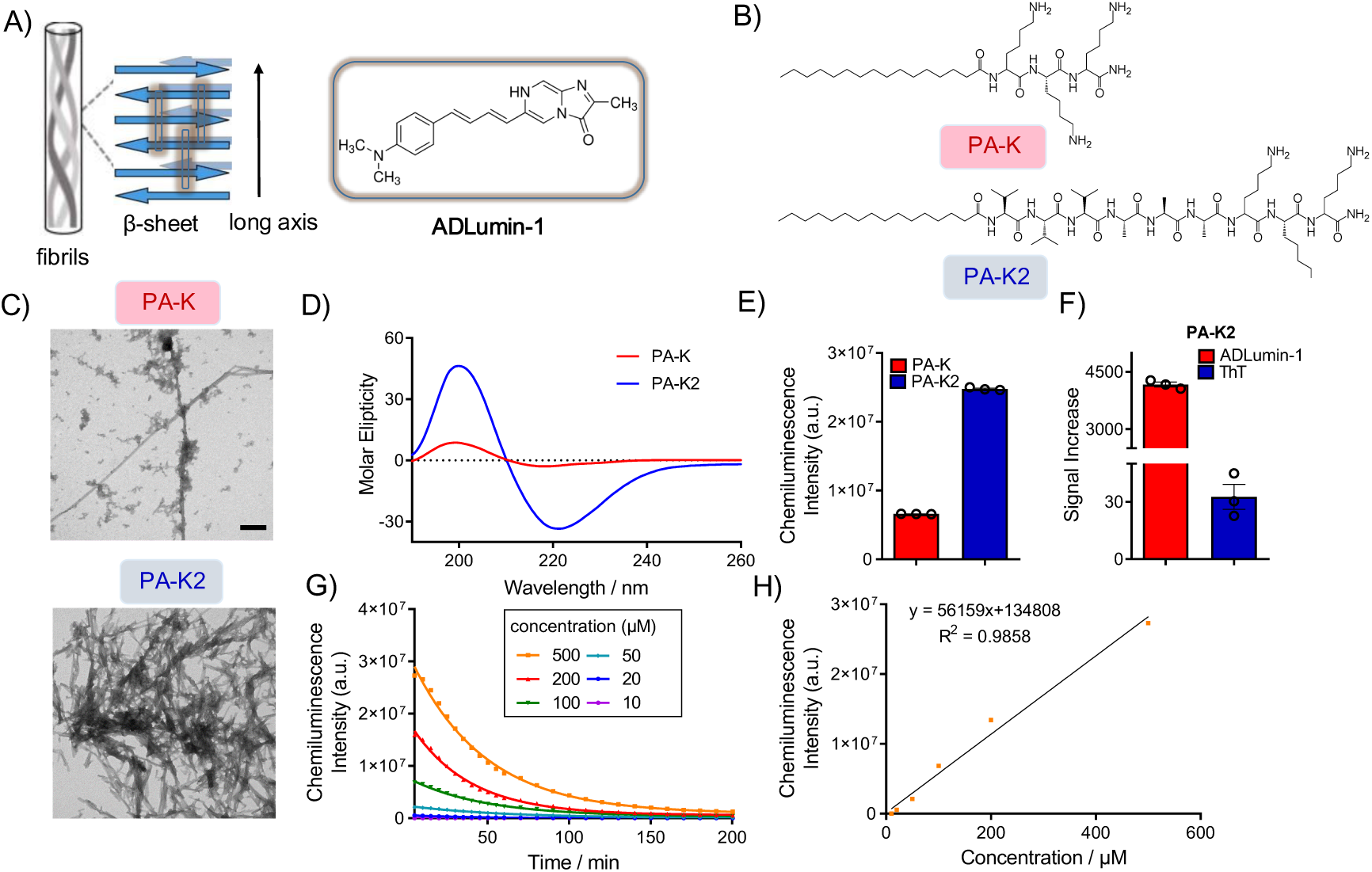
ADLumin-1’s specificity to β-sheet-rich structures. (**A**) Chemical structure of ADLumin-1 and mechanism for sensing β-sheet-rich structure. (**B**) Chemical structure of PA-K/K2. (**C**) Representative TEM images for PA-K/K2 (500 μM). (**D**) CD spectra of PA-K/K2 (10 μM). PA-K2 showed typical peaks indicating the formation of β-sheet-rich structure, while PA-K showed a low degree of β-sheet. (**E**) Chemiluminescence intensity of ADLumin-1 with PA-K or PA-K2 (500 μM). (**F**) SNR of ADLumin-1 and ThT with PA-K2 (500 μM). (**G**) Chemiluminescence signal profile of ADLumin-1 with different concentrations of PA-K2 (10-500 μM) from 5 to 200 min. (**H**) Linear regression of the highest signal of ADLumin-1 with different concentrations of PA-K2 (10-500 μM).

After incubating 500 μM model peptides in PBS buffer (pH = 7.4) for 72 h with constant shaking, the protein misfolding was studied by transmission electron microscopy (TEM). As shown in Fig. 1C and Fig. S2B, abundant fibrillar structures can be observed in the PA-K2 and PA-E2 groups, while PA-K and PA-E groups showed a small amount of fibrillar structure. To further investigate the secondary structure of the four model peptides, circular dichroism (CD) spectra were performed. As shown in Fig. 1D and Fig. S2C, PA-K2, and PA-E2 were predominantly β-sheets, evidenced by a typical negative band around 220 nm and a positive band around 200 nm. In contrast, PA-K contained a low degree of β-sheet structure, and PA-E was composed of random coils.

To study the detecting capability of ADLumin-1 for β-sheet fibrils, we first incubated ADLumin-1 (15 µM) with PA-K or PA-K2 (500 μM) in PBS buffer. Compared to the background signal, the chemiluminescence signal was dramatically increased in the PA-K2 group (signal increase fold = 4168.25 ± 88.00), while there was a moderate increase in the PA-K group (signal increase fold = 248.10 ± 19.95) (Fig. 1E). Remarkably, the intensity ratio of PA-K2/PA-K was 16.8-fold, suggesting that ADLumin-1 has excellent specificity towards β-sheet-rich fibrils. Next, we compared the signal increase of ADLumin-1 to the gold standard dye ThT, and found that ADLumin-1 showed a 127.73-fold higher SNR compared with fluorescence dye ThT when interacting with PA-K2 (Fig. 1F). In addition, we found that the intensities of PA-K2 were concentration-dependent and ADLumin-1 provided a linear response with 10-500 μM of PA-K2 (Fig. 1G, H).

Next, we studied the chemiluminescence response of ADLumin-1 with PA-E/PA-E2. Similar to the results of PA-K/PA-K2, ADLumin-1 showed a selective signal increase to PA-E2 over PA-E (Fig. S2D), and ADLumin-1 showed a 52.02-fold higher SNR compared with ThT when interacting with PA-E2 (Fig. S2E). As shown in Fig. S2F and Fig. S2G, ADLumin-1 showed increased signal with linear responses in the presence of different concentrations of PA-E2 (1-100 μM). Interestingly, the model peptides showed different decay profiles. PA-K2 showed a half-life of 31.08 min, while PA-E2 displayed a half-life of 42.43 min. This may be attributed to the different turnover rates of ADLumin-1 upon binding with the hydrophobic tunnels of the β-sheet fibrils (*42*). In addition, we also tested the interaction of ADLumin-1 with α-helix-rich human serum albumin (HSA), and compared the signal difference of HSA and PA-K2. Interestingly, ADLumin-1 showed very fast signal decay when interacting with HSA, and the chemiluminescence intensity of PA-K2 is 105.87-fold higher than HSA after 30 min incubation (Fig. S2H, I). The slow decay in the presence of β-sheet-rich structure indicates that ADLumin-1 forms a strong and specific bond with the hydrophobic tunnel within the β-sheet-rich structure, whereas its interaction with albumin appears to be transient and non-specific.

At the early stage of protein misfolding, the monomer unit of misfoldons assemble to form oligomers, and this process contributes to misfoldon toxicity (*5*). However, ThT has limited responsive capacity for oligomers (*51*). To further investigate the β-sheet-responsive capability of ADLumin-1 to oligomers, we used the CATCH peptide pair as the model. The CATCH pairs contain CATCH+ (QQKFKFKFKQQ) and CATCH-(EQEFEFEFEQE) peptides that immediately assemble upon mixing to form oligomers (*52, 53*). The peptide structure is shown in Fig. S3A, and the successful formation of β-sheet-rich oligomers was confirmed by TEM (Fig. S3B) and CD spectrum (Fig. S3C). As we expected, ADLumin-1 showed a significantly increased chemiluminescence signal when interacting with the CATCH peptide pair (Fig. S3D, E). The SNR of ADLumin-1 detection is 1.67-fold higher compared to ThT (Fig. S3D).

Collectively, these results suggested that ADLumin-1 has excellent chemiluminescence response and selectivity for β-sheet-rich fibrils.

### ADLumin-1 is a generic chemiluminescence probe for misfoldons

To investigate the interaction of ADLumin-1 with misfoldons, we prepared fibrils of α-syn, Aβ, tau, and TDP-43 proteins, which were confirmed by TEM imaging and ThT fluorescence assay (Fig. S4). Upon mixing with the misfoldons, the chemiluminescence signals of ADLumin-1 considerably amplified, showing significantly enhanced signal increases of 111.8 ± 13.1-fold for α-syn, 170.6 ± 23.3-fold for Aβ, 26.2 ± 4.1-fold for tau, 14.7 ± 1.4-fold for TDP-43. Compared to ThT, the increased folds were much larger, and the largest difference was 82.8 ± 9.7-times for α-syn, 74.5 ± 10.2-times for Aβ, 14.9 ± 2.3-times for tau, 11.4 ± 1.1-times for TDP-43 (Fig. 2A). The signal to background ratio of ADLumin-1 to ThT when interacting with misfolded tau or TDP-43 were 11.88 ± 1.55 and 11.50 ± 0.87, respectively (Fig. S5A). The results strongly suggested that ADLumin-1 has much better sensitivity for detecting misfoldons than the golden standard ThT. We also found ADLumin-1 showed SNR of 222.15 ± 29.71 at 180 min after mixing with misfolded α-syn (Fig. S5B), the long chemiluminescence duration time indicated that ADLumin-1 is favorable for in vitro solution test. In addition, with an increased time of shaking, the probe showed gradually increased signals of various misfolded proteins from 12-72 h (Fig. S5C). Furthermore, we compared the chemiluminescence signal of ADLumin-1 when interacting with fibrillar or monomeric forms of misfolded proteins. As shown in Fig. S5D, ADLumin-1 showed significantly higher signals when interacting with fibrils compared to monomers, which was 26.38 ± 3.09-times for α-syn, 22.75 ± 1.36-times for Aβ, 2.83 ± 0.37-times for tau, 6.54 ± 0.49-times for TDP-43. We also tested the signal decay profile of ADLumin-1 with various misfoldons. Interestingly, different misfoldons showed different chemiluminescence signal decay half-life times (Table S1). This indicated ADLumin-1 could not only detect but also potentially differentiate the misfoldons. Next, we performed molecular docking of ADLumin-1 with the Cryo-EM structures of α-syn, Aβ, tau, and TDP-43. As shown in Fig. 2B and Table S2, ADLumin-1 could bind along the long axis of the fibrils of those misfoldons by interacting with the hydrophobic tunnels that are formed by Ala, Val or Phe (Fig. 2B, C). To further confirm the detecting capability of ADLumin-1 to various misfoldons, fibrils of prion, amylin, and transthyretin were prepared and the Cryo-EM structures of prion, amylin, and transthyretin proteins were used for molecular docking. ADLumin-1 showed higher SNR compared to ThT when interacting with those misfoldons (Fig. S6A), and displayed a similar binding position along the long axis of those β-sheet structures and hydrophobic interaction with Val and Phe (Fig. S6B).

**Fig. 2.**
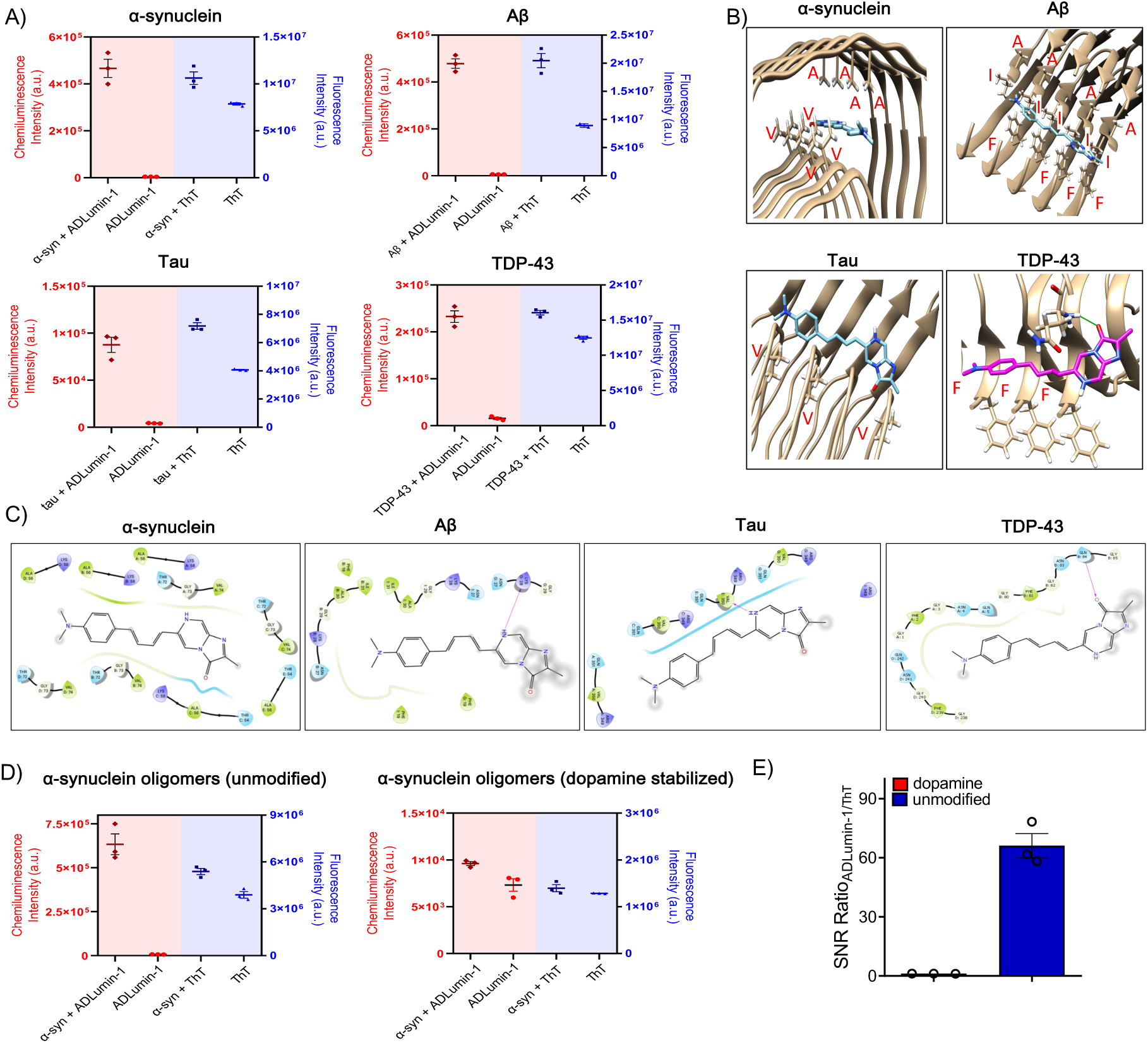
ADLumin-1 is a generic chemiluminescence probe for misfoldons. (**A**) The chemiluminescence or fluorescence signal of ADLumin-1 or ThT (final concentration: 15 μM in 5% DMSO/PBS buffer) when mixing with misfolded proteins (misfoldons). ADLumin-1 showed significantly higher signal increases compared to ThT. (**B**) Molecular docking of ADLumin-1 with misfoldons. ADLumin-1 showed similar binding poses that bind along the long axis of several misfoldons. (**C**) Key interacting residues of ADLumin-1 to misfoldons, ADLumin-1 interacts with the hydrophobic tunnels that are formed by Ala, Val, Ile or Phe. (**D)** The chemiluminescence or fluorescence signal of ADLumin-1 or ThT (final concentration: 15 μM in 5% DMSO/PBS buffer), and SNR ratio of ADLumin-1 to ThT when mixing with unmodified α-syn oligomers or dopamine-stabilized α-syn oligomers. ADLumin-1 showed stronger signal enhancements and higher SNR compared to ThT when mixing with β-sheet-rich unmodified α-syn oligomers.

To investigate the interaction of ADLumin-1 with various fibril strains/polymorphs, we prepared several fibrils with or without salt. It has been reported that salt ions could change the aggregation propensity and generate different fibrillar structures (*54, 55*). In the presence of salt, Aβ fibrils are associated with smaller diameters (*54*), α-syn proteins tend to form short fibrils (*56*), tau proteins could rapidly oligomerize and form diverse fibrillar structures (*55*), and TDP-43 proteins tend to aggregate with higher turbidity (*57*). The fibril morphologies were confirmed by TEM imaging (Fig. S7A). We tested the chemiluminescence signal when mixing the samples with ADLumin-1 (Fig. S7B). Interestingly, ADLumin-1 showed distinct signal decay profiles when interacting with different fibril strains/polymorphs. The different responses may result from different binding modes of ADLumin-1 to the fibrils. We further conducted the docking experiments with various human-derived or recombinant fibril polymorphs (*58–61*). As shown in Fig. S8 and Table S3, the probe binds to all examined fibrils, frequently interacting with hydrophobic residues such as Ala, Val, Phe, and Tyr. Notably, ADLumin-1 generally binds along the long axis of most recombinant fibrils (PDB IDs for Aβ: 5OQV, 2LMO, 2NAO; α-syn: 6PEO, 6A6B, 6SSX; TDP-43: 8QX9, 8QXA, 8QXB) as well as human-derived fibrils (PDB IDs for α-syn: 6XYO, 6XYQ; TDP-43: 7PY2, 8CG3). Interestingly, we also note that human-derived fibrils (PDB IDs for Aβ: 2M4J, 8QN6, 8QN7; α-syn: 8A9L; TDP-43: 8CGG) exhibit more complex binding poses, indicating the greater heterogeneity of fibrils in human brains. For tau fibrils, ADLumin-1 shows distinct binding poses across different fibrils, likely due to the diverse isoforms and morphologies of tau fibrils (*58*).

To study the detecting capability of ADLumin-1 with oligomer species, we used ADLumin-1 to detect several purified protein oligomers and used ThT for comparison. The TEM and SEC characterization of oligomers were shown in Fig. S9A, B. While ThT showed slightly changed signals, the signal enhancement of ADLumin-1 with oligomers was much larger, which was 42.38 ± 3.30-fold for Aβ, 25.00 ± 0.59-fold for tau, and 17.17 ± 3.58-fold for TDP-43 (Fig. S10A). The SNR ratio of ADLumin-1 to ThT was 34.50 ± 2.69-fold, 23.81 ± 0.56-fold, and 16.46 ± 3.43-fold for Aβ, tau and TDP-43 oligomers, respectively (Fig. S10B).

To further investigate the β-sheet-responsive capability of ADLumin-1 to oligomers, we used ADLumin-1 to detect dopamine-induced α-syn oligomers and unmodified α-syn oligomers and used ThT for comparison (*62, 63*). As TEM imaging results showed, dopamine-induced α-syn oligomers showed disordered structures while unmodified α-syn oligomers showed β-sheet-rich structures (Fig. S9A), which were in accordance with previous literature (*62*). As shown in Fig. 2D, ThT showed negligible signal change for dopamine-induced α-syn oligomers but slightly increased signals with β-sheet-rich unmodified oligomers. The signal enhancement for ADLumin-1 was 91.59 ± 12.08-fold for unmodified α-syn oligomers and 1.31 ± 0.04-fold for dopamine-induced α-syn oligomers (Fig. 2D). The SNR ratio of ADLumin-1 to ThT was 66.14 ± 8.72-fold and 1.21 ± 0.04-fold for unmodified α-syn oligomers and dopamine-induced α-syn oligomers, respectively (Fig. 2D), which further indicated the interaction of ADLumin-1 to β-sheet-rich structures.

Taken together, our results suggested that ADLumin-1 could serve as a generic chemiluminescence probe for detecting misfolded β-sheet structures.

### ADLumin-1 for visualizing misfolded tau and TDP-43 in vitro and in vivo

Our in vitro solution studies indicated that ADLumin-1 is a generic imaging probe for various misfoldons, and we previously demonstrated that ADLumin-1 could be used as an in vivo imaging probe for Aβ in transgenic mouse models (*42*). To further expand the scope of in vivo imaging for misfoldons, we investigated whether ADLumin-1 could non-invasively visualize other misfoldons that serve as key biomarkers in neurodegenerative diseases.

The aggregation of tau protein is a crucial factor that drives AD and is also closely associated with Pick disease, progressive supranuclear palsy and other tauopathies (*12*). In Fig. 2A, we showed that the chemiluminescence of ADLumin-1 could be enhanced in the presence of tau aggregates. To assess the binding affinity of ADLumin-1 with tau aggregates, we measured the binding constant, which showed moderate binding with K*d* of 2.18 μM (Fig. 3A). Next, we used ADLumin-1 to stain AD patient brain slides, and found that ADLumin-1 was capable of visualizing tau tangles, confirmed by subsequent anti-p-tau antibody staining (Fig. 3B). Furthermore, we conducted colocalization analysis of ADLumin-1 and antibody staining group by using Scatter J. ADLumin-1 can label most of the tau tangles and colocalized with antibody staining, showing Pearson’s R values of 0.88 (Fig. S10C). To validate ADLumin-1’s capacity for in vivo tau imaging, we used 10-month-old P301L mice. Indeed, ADLumin-1 provided considerable large differences between P301L and WT mice at different time points, and the largest margin was about 3.0-fold at 360 minutes after i.p. injection of the probe (Fig. 3C, D). Taken together, our data demonstrated that ADLumin-1 could detect tau tangles in vivo.

**Fig. 3.**
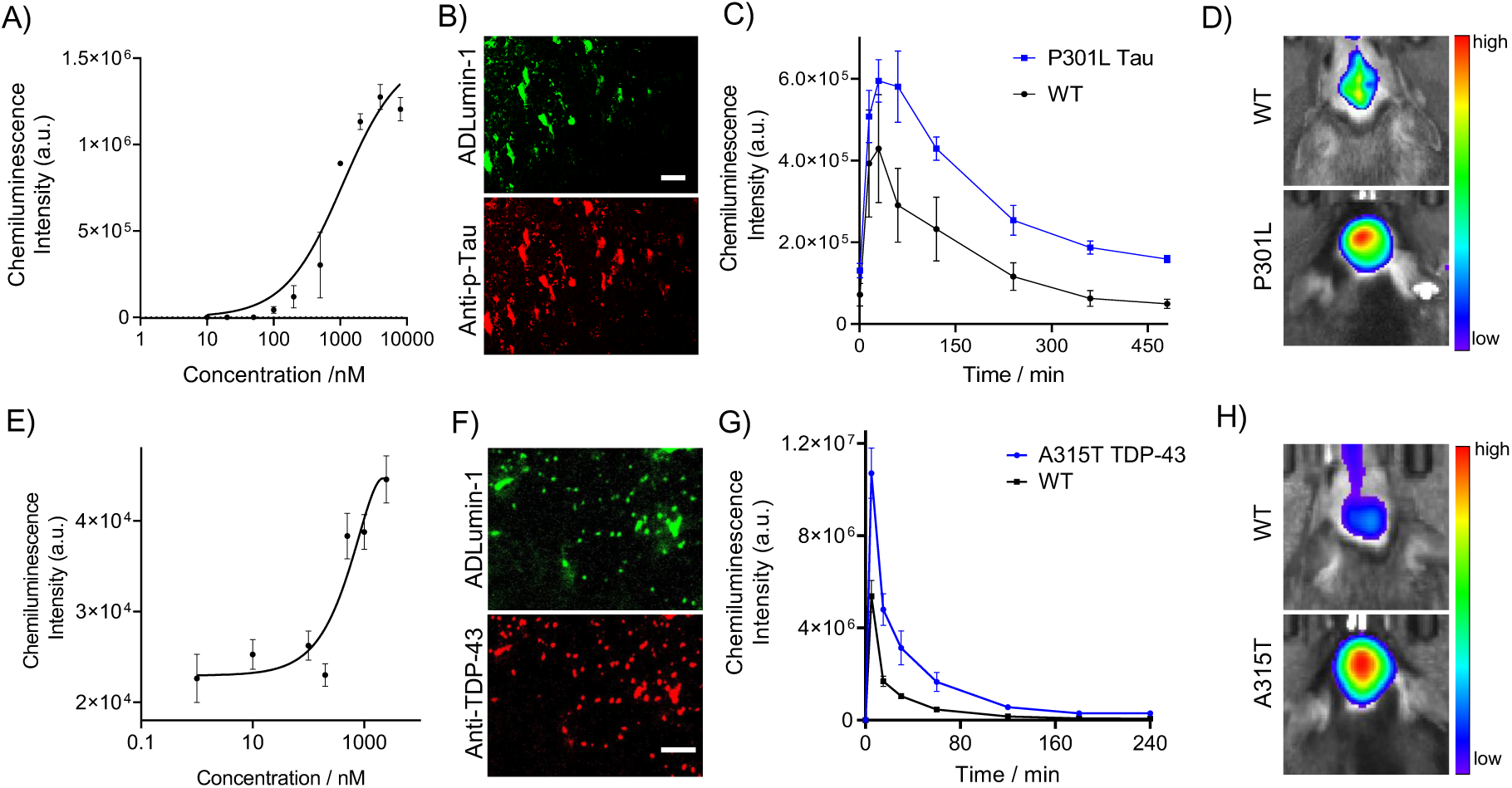
ADLumin-1 as a generic chemiluminescence probe for tau and TDP-43 aggregates in vitro and in vivo. (**A**) Binding constant measurement of ADLumin-1 with tau aggregates (K*d* = 2.18 μM). (**B**) Pathological tau tangle staining in AD patient brain slides (female, 95 years old) with ADLumin-1, the labeling of tau tangle was confirmed by subsequent staining of anti-p-tau antibody. (**C**) Time courses of chemiluminescence signals from ADLumin-1 after i.p. injection with P301L mice and WT mice (female, 10-month-old, n = 5). (**D**) Representative brain images of P301L and WT mice at 60 minutes post-injection. (**E**) Binding constant measurement of ADLumin-1 with TDP-43 aggregates (K*d* = 2.34 μM). (**F**) Pathological TDP-43 aggregates staining in A315T transgenic mice (female, 6-month-old) with ADLumin-1, the labeling of TDP-43 deposit was confirmed by subsequent staining of anti-TDP-43 antibody. (**G**) Time courses of chemiluminescence signals from ADLumin-1 after i.v. injection with A315T mice and WT mice (female, 6-month-old, n =5). (**H**) Representative brain images of A315T and WT mice at 60 minutes post-injection.

TDP-43 is a protein that plays key roles in regulating gene expression and RNA processing in cells (*64–66*). In amyotrophic lateral sclerosis (ALS), TDP-43 forms abnormal aggregates in the motor neurons of the brain and spinal cord, leading to dysfunction and eventual degeneration (*64*). Consequently, there has been considerable interest in developing drugs that target TDP-43 aggregation as a potential therapeutic strategy for ALS (*64, 67*). Our in vitro solution results showed there was a 14.7-fold increase in chemiluminescence intensity upon binding to TDP-43 aggregates, compared to the control group (without TDP-43) (Fig. 2A). In line with this result, we further measured the dissociation constant with the TDP-43 aggregates, showing K*d* of 2.34 μM (Fig. 3E). To validate whether ADLumin-1 could stain TDP-43 deposits in brain slides, we used a 6-month-old A315T mouse brain slides and stained with ADLumin-1. As we expected, ADLumin-1 provided clear highlights of the TDP-43 deposits, confirmed by subsequent staining with anti-TDP-43 antibody (Fig. 3F). Furthermore, we conducted colocalization analysis of ADLumin-1 and antibody staining group by using Scatter J, which showed colocalization with Pearson’s R values of 0.81 (Fig. S10D). Lastly, we also demonstrated that ADLumin-1 could be used to image TDP-43 in vivo with transgenic ALS mice (A315T) and found that the difference between Tg/WT was about 3.59-fold (Fig. 3G, H). However, we found that i.v. injection of ADLumin-1 was necessary to obtain the differences in imaging signals. This is different from the cases of mouse models of Aβ, tau, and α-syn, with which i.p. injection could be used for imaging. It is unclear what factors are causing this difference and may be due to the micro-environment of the peritoneum of A315T mice being different from other mouse models.

Taken together, our data indicated that ADLumin-1 was capable of targeting tau and TDP-43 both in vitro and in vivo and could be used to monitor cerebral tau and TDP-43 misfolding in transgenic mouse models.

### ADLumin-1 enables highly sensitive and selective detection of misfolded α-syn in vitro

To further study the detecting capability of ADLumin-1, we selected α-syn as the model misfoldon. α-syn is a major component of lewy bodies/neurites and was widely considered a typical pathological hallmark of Parkinson’s disease (PD) and Lewy body dementia (LBD) (*13*). Currently, there is no disease-modifying cure for PD (*5*). A few fluorescence probes can detect α-syn fibrils in vitro (*68, 69*), and exploratory antibody-based probes can detect α-syn fibrils in vivo (*70*). To date, there is still a significant need for effective small molecular optical imaging probes for both in vitro and in vivo studies targeting α-syn aggregates(*71*).

The above data (Fig. 2) indicate that ADLumin-1 could be used as a chemiluminescence probe to detect α-syn fibrils. First, we measured the binding affinity of ADLumin-1 to α-syn fibrils. Through titration of ADLumin-1, the dissociate constant (K*d*) was calculated as 0.76 µM (Fig. 4A). With increased time of shaking, ADLumin-1 showed increased signals with α-syn from 12-72 h and the chemiluminescence decay spectra were shown in Fig. 4B. The emission spectra of ThT with α-syn aggregation from 12-72 h were also shown in Fig. S11A. To further confirm the detecting capability of ADLumin-1 to misfolded β-sheet structure, two inhibitors of Apigenin and Resveratrol for α-syn were used for inhibiting α-syn aggregation (*72, 73*). We conducted sedimental experiments by following previous protocols (*74*). Briefly, we prepared α-syn samples with or without Apigenin and Resveratrol and shook them for 0, 12, 24, 48, and 72 h. The samples at the final time point (72 h) were used for TEM imaging and samples at each time point were subjected to centrifugation to separate soluble and insoluble α-syn, followed by detecting the soluble species with gel electrophoresis and detecting the insoluble pellets with ADLumin-1 (workflow in Fig. S12A). The TEM images captured at the end of incubation revealed fibrils in α-syn group but soluble species in the inhibitor-treated group (Fig. S12B). The α-syn only group showed increased α-syn fibril formation from 0 to 72 h, evidenced by enhanced chemiluminescence intensity (Fig. 4C) and decreased remaining soluble species detected by gel electrophoresis (Fig. S12C, D). Apigenin and Resveratrol groups showed slightly changed ADLumin-1’s signal (Fig. 4C) and soluble species (Fig. S12C-E). These data indicated that inhibitors significantly inhibit the formation of α-syn fibrils, and this process can be detected by ADLumin-1. In addition, we examined the lower limit of detection (LLOD) of ADLumin-1 to α-syn fibrils. We spiked 0-10 pg/ml (0, 70, 140, 350, 700 fM) α-syn fibrils in PBS buffer and added ADLumin-1 for detection. As shown in Fig. 4D, ADLumin-1 can detect 1 pg/ml α-syn fibrils in PBS buffer, showing an SNR of 2.31 ± 0.39. Whereas no significant fluorescence difference could be observed with ThT (Fig. S11B), suggesting ADLumin-1 has a better detection limitation than ThT. In addition, ADLumin-1 showed a linear response with 1-10 pg/ml (0-700 fM) of α-syn fibrils.

**Fig. 4.**
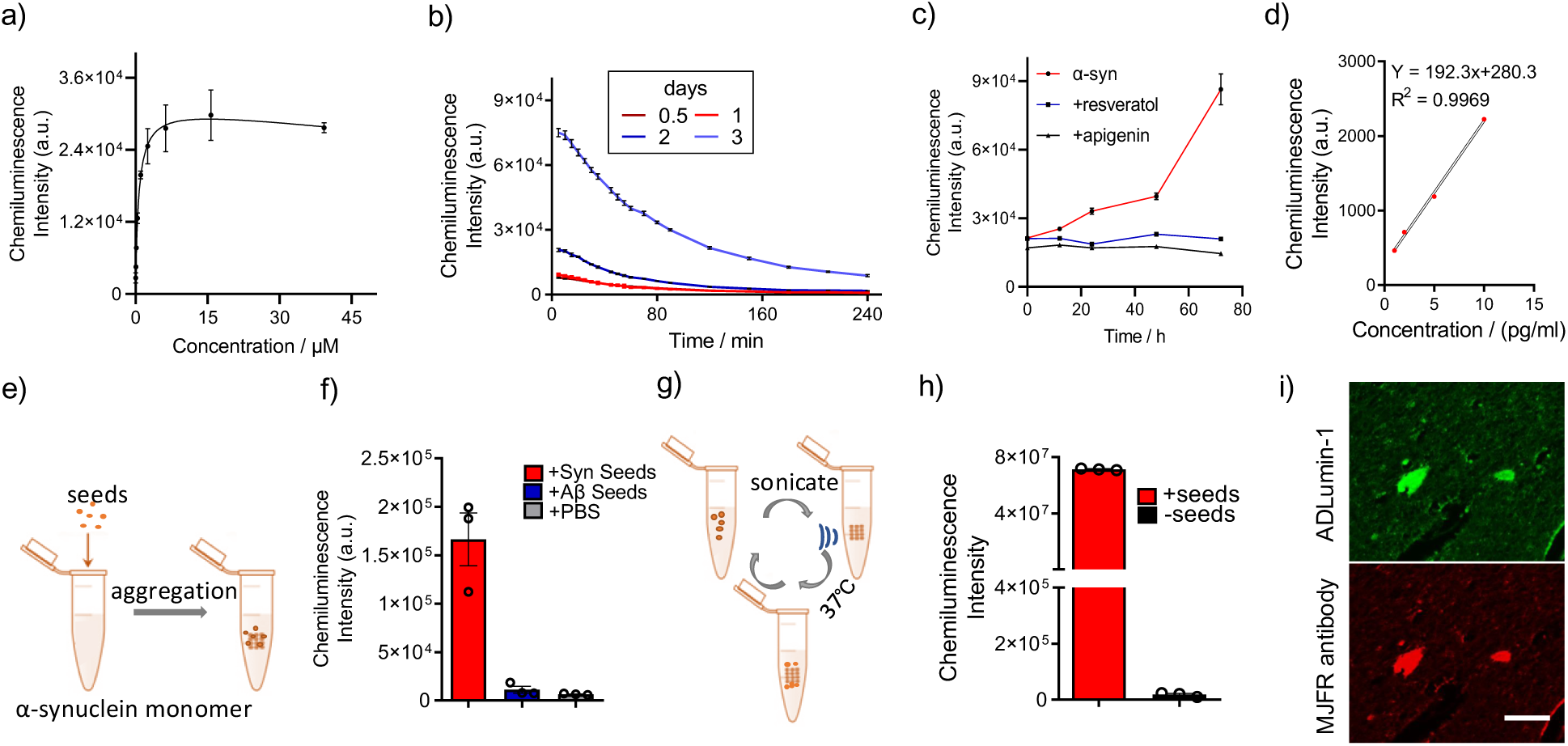
ADLumin-1 enables highly sensitive and selective detection of misfolded α-syn in vitro. (**A**) Binding affinity assay of ADLumin-1 (0-40 μM) with 250 nM α-syn fibrils (K*d* = 0.76 µM). (**B**) Signal decay profile of α-syn with increased time of aggregation (12-72 h) by using ADLumin-1. (**C**) Chemiluminescence intensity of ADLumin-1 with insoluble α-syn samples in sedimentation experiments. α-syn proteins showed increased signal during 12-72 h. With the inhibitors (Apigenin or Resveratrol), the signal showed negligible changes over time, indicating the inhibitors significantly inhibit the formation of α-syn fibrils, and this process can be detected by ADLumin-1. (**D**) Direct detection of α-syn fibrils in PBS buffer with ADLumin-1, which indicates a lower limit of detection at 1 pg/mL (70 fM). (**E**) Diagram of seeding experiments. α-syn monomers in PBS solution (200 μL, 25 μM) were added with 20 μL α-syn seeds (2.5 μM, final concentration), Aβ seeds (2.5 μM, final concentration), or PBS and incubated for 48 h. (**F**) Chemiluminescence intensity of different seeding groups. ADLumin-1 only showed significant signal amplification in the α-syn seeds group but negligible signal in the Aβ seeds and PBS group. (**G**) Diagram of protein misfolding cyclic amplification (PMCA) workflow. (**H**) Chemiluminescence intensity of α-syn monomer with or without α-syn seeds (1 pg/ml, spiked in CSF). ADLumin-1 enables selective detection of 70 fM α-syn aggregates by combining with PMCA technology. (**I**) Pathological α-syn aggregates staining in PD patient brain slides (temporal lobe, male, 85 years old) with ADLumin-1. The labeling of α-syn aggregates was confirmed by subsequent staining of anti-α-syn aggregate antibody.

Since the increased level of misfolded α-syn in the cerebrospinal fluid (CSF) has been closely associated with PD pathology (*36, 75*), we explored the feasibility of detecting α-syn fibrils in CSF with ADLumin-1. If this feasibility is validated, ADLumin-1 can be potentially used as an agent for in vitro diagnosis of PD and LBD. To this end, we first spiked 70 fM α-syn fibrils into the CSF and then used ADLumin-1 to detect the signal increase. However, ADLumin-1 failed to detect the signal increase, possibly due to the interference of the non-specific binding in CSF. To solve this problem, we utilized the strong seeding property of α-syn fibrils for signal amplification. Recent evidence showed a strong seeding effect of α-syn, where pre-formed misfolded α-syn (seeds) can accelerate the aggregation of α-syn monomers (*13, 76*). To verify whether ADLumin-1 can be used to monitor the seeding effect, we first prepared α-syn seeds and added them into a solution of α-syn monomers, followed by 48 h of constant shaking (Fig. 4E). Aβ aggregates were also prepared as the seeds for validating the selectivity of our method. As shown in Fig. 4F, ADLumin-1 only showed significant signal turn-on in the α-syn seeding group, but not in the control group. This result inspired us to perform further signal amplification in CSF by combing ADLumin-1 with protein-misfolding cyclic amplification (PMCA), which utilized the seeding effect with cyclic sonication for ultrasensitive detection of α-syn (Fig. 4G) (*35, 36*). As shown in Fig. 4H, ADLumin-1 successfully identified 70 fM α-syn aggregates in CSF, showing 3955.67 ± 32.14-fold increase of the signal in the presence of α-syn seeds compared to the control group. Furthermore, we compared the detecting capability of ADLumin-1 with cyclic amplification by using a microplate reader, and used ThT for comparison (Fig. S13A). By comparing the highest signal of ADLumin-1 and ThT for 35 pM α-syn aggregates in CSF, we found ADLumin-1 showed 32.56 ± 5.79-fold higher signal compared to ThT (Fig. S13B). These data indicated that ADLumin-1 could be applied for ultra-sensitive detection of misfolded α-syn in solutions and CSF, and has the potential for in vitro diagnosis via detecting α-syn fibrils in CSF.

To study the binding of ADLumin-1 to pathological α-syn aggregates in the brain, we stained brain tissues from a Parkinson’s disease patient (temporal lobe, male, 85 years old) with ADLumin-1(*77*). To confirm the labeling of ADLumin-1 to α-syn aggregates, the brain tissues were further stained with an anti-α-syn antibody. As shown in Fig. 4I, ADLumin-1 successfully labeled Lewy bodies and Lewy neurites from the patients’ brains with strong fluorescence (Note: ADLumin-1 itself is also fluorescent). This result suggested that ADLumin-1 can be a complemental agent for postmortem brain tissue staining to characterize synuclein pathologies.

### In vivo detection of α-syn fibrils with ADLumin-1

To investigate the in vivo detection of cerebral α-syn pathology, we intracranially injected α-syn aggregates (62.5 μM, 400 nL) into the substantial nigra of C57/BL6J mice (n = 5, female, 22-month-old). After recovering for one month, the mice were subjected to in vivo 3D diffuse luminescent imaging tomography (DLIT) with ADLumin-1 (Fig. 5A). As shown in Fig. 5B, ADLumin-1 specifically labeled α-syn aggregates in the deep brain and generated chemiluminescence signals. The reconstructed images also suggested that ADLumin-1 is highly selective for α-syn aggregates and has no apparent binding to non-α-syn proteins in the brain.

**Fig. 5.**
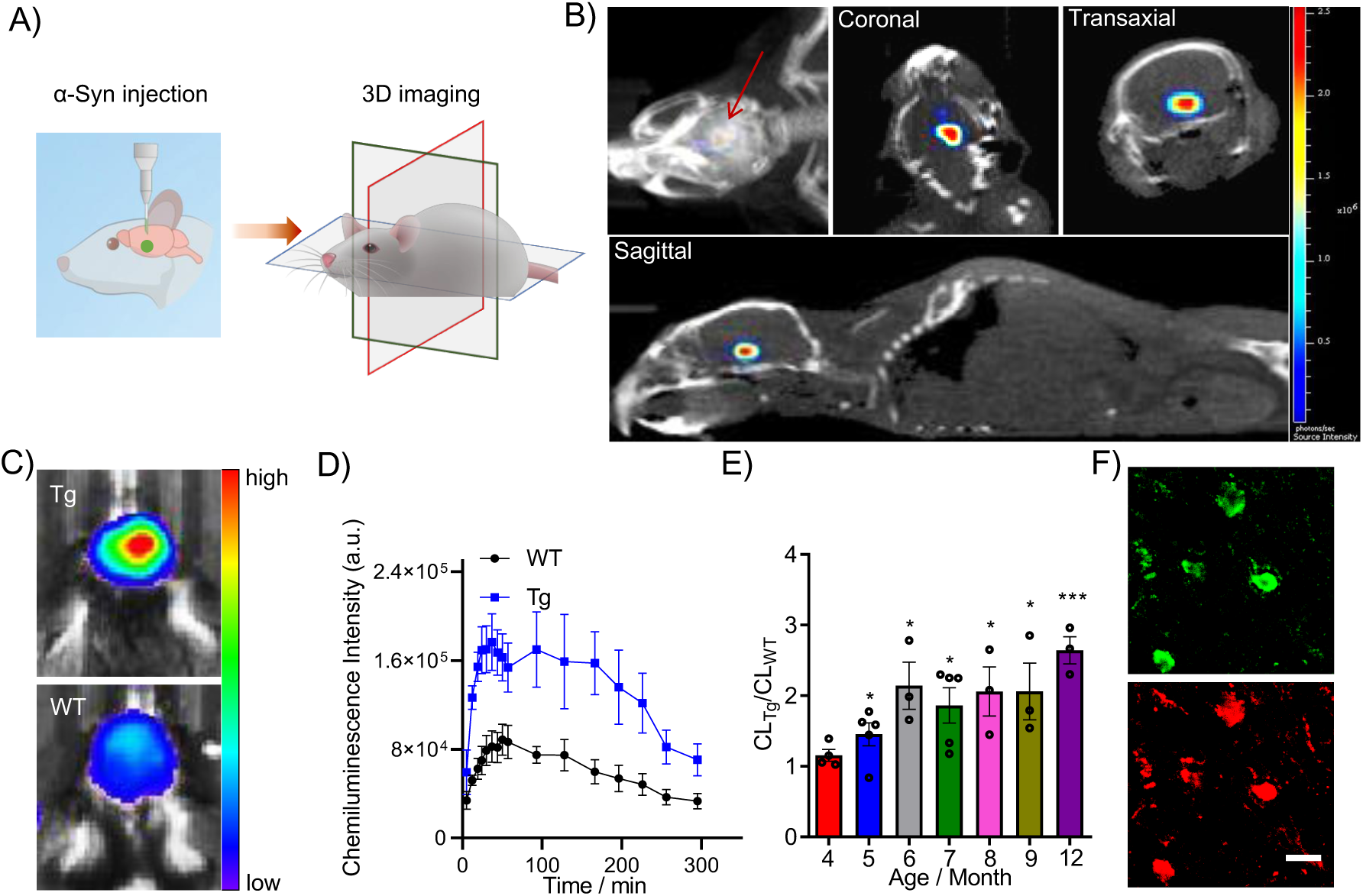
In vivo detection of α-syn fibrils with ADLumin-1. (**A**, **B**) Illustration diagram (**A**) and representative 3D DLIT images (**B**) of C57/BL6J mice that were intracranially injected with α-syn aggregates (62.5 μM, 400 nL) into the substantial nigra region of the brain. ADLumin-1 showed selective binding and signal amplification to the cerebral α-syn aggregates in vivo. (**C**, **D**) Representative images (**C**) and quantitative signal (**D**) of in vivo imaging of transgenic mice and wild-type mice (6-month-old, female, n ≥ 3) after i.p. injected with ADLumin-1 (8 mg/kg). Images were acquired at 30 min postinjection. (**E**) Monitor the progression of α-syn aggregation with ADLumin-1 (8 mg/kg) in transgenic A53T mice (female, 4 to 12-month-old, n ≥ 3). The ratio of chemiluminescence intensity of transgenic mice and wild-type mice increased with disease progression. (**F**) Ex vivo histological observation of A53T transgenic mice and wild-type mice. A53T mice showed α-syn aggregates labeled by ADLumin-1, which was confirmed by subsequent staining of an antibody that is specific to α-syn aggregates (scale bar = 20 μm).

To further study the in vivo detecting capability of ADLumin-1, transgenic A53T mice (n = 5, female) and wild-type control (n = 5, female) were subjected to in vivo imaging at different ages (4 to 12 months old). The A53T mice overexpress human α-syn with a PD-associated mutation (A53T) driven by the prion promoter, and exhibit abnormal accumulation and aggregation of α-syn proteins in the brain (*78*). After i.p. injection of ADLumin-1, transgenic mice (Tg) showed higher signals from the brain areas, compared with the wild-type (WT) mice (Fig. 5C,D). As shown in Fig. 5D, the signal peaked around 40 min after the probe injection for A53T and WT mice, showing a 2.14-fold signal difference in the brain area. The chemiluminescence signal of ADLumin-1 exhibited a long duration time in transgenic mice after i.p. injection, which showed SNR of 177.30 ± 75.38 at 180 min postinjection (Fig. S13C). Notably, ADLumin-1 was able to detect the difference between A53T and WT mice at the age of 4 months old, and the difference was about 1.28-fold (Fig. 5E). Most importantly, ADLumin-1 enabled longitudinal monitoring of α-syn accumulation, as evidenced by the increasing signal ratios between A53T/WT mice with age (Fig. 5E). The ratio reached 2.64-fold for the 12-month-old mice. To further confirm the binding of ADLumin-1 to α-syn inclusion in vivo, we performed ex vivo histology staining after the imaging of the mice at the age of 12 months old. The mice brains were harvested and cut into slides, followed by co-staining with anti-α-syn antibody. As we expected, the ex vivo histological staining indicated strong labeling of diffuse cytoplasmic asyn and Lewy bodies (Fig. 5F) in the transgenic mice brain. These data further suggested that ADLumin-1 can penetrate mice brains and differentiate transgenic mice from wild-type mice.

### Screening of NIR fluorophores for selective bio-orthogonal-ChRET imaging of misfolded α-syn

Despite ADLumin-1 enabling in vivo chemiluminescence imaging of cerebral misfoldons, it still exhibits two limitations that require further attention: 1) its short emission could lead to significant absorption in biological tissues, which hampers deep penetration of the photons for in vivo imaging; 2) ADLumin-1 lacks selectivity to misfolded α-syn.

To overcome the limitations, we reasoned that bio-orthogonal ChRET (chemiluminescence resonance energy transfer) imaging could be utilized. In recent years, bio-orthogonality has been widely applied for in vitro and in vivo imaging (*79*). While click-chemistry has been routinely used to achieve bio-orthogonality via forming covalent bonds, the non-conjugated pairing could also enable bio-orthogonality (*79*). Recently, our group demonstrated that the non-conjugated pairing of two imaging probes could be used for bio-orthogonal imaging of Aβ aggregates via <u>d</u>ually-<u>a</u>mplify <u>s</u>ignal via ChRET (DAS-ChRET) (*42*). The non-conjugated pairing strategy utilized the proximity of two imaging probes (one chemiluminescence probe, and one near-infrared fluorescence probe) that gave a longer emission wavelength for a better SNR. Theoretically, for such bio-orthogonal-ChRET, the probe pairs need to meet the following requirements (Fig. 6A, A and B sites): 1) a chemiluminescence imaging probe (donor) can bind at site A and generate a strong chemiluminescence signal; 2) a near-infrared (NIR) fluorophore (acceptor) can bind at site B and generate strong NIR fluorescence signals; 3) the distance between site A and site B is less than 10 nm; 4) the emission wavelength of the chemiluminescence probe should be overlapped with the excitation wavelength of the fluorophore. As the misfoldons share common features of the abundant β-sheet-rich structures that offer multiple binding sites, we hypothesized that ADLumin-1 could be a generic probe to target the long axis of laminated cross-β architecture and serve as a donor probe. By applying a selective NIR fluorescence probe for misfolded α-syn, the bio-orthogonal-ChRET strategy could be used for selective imaging of α-syn fibrils due to the rigorous proximity requirement (less than 10 nm).

**Fig. 6.**
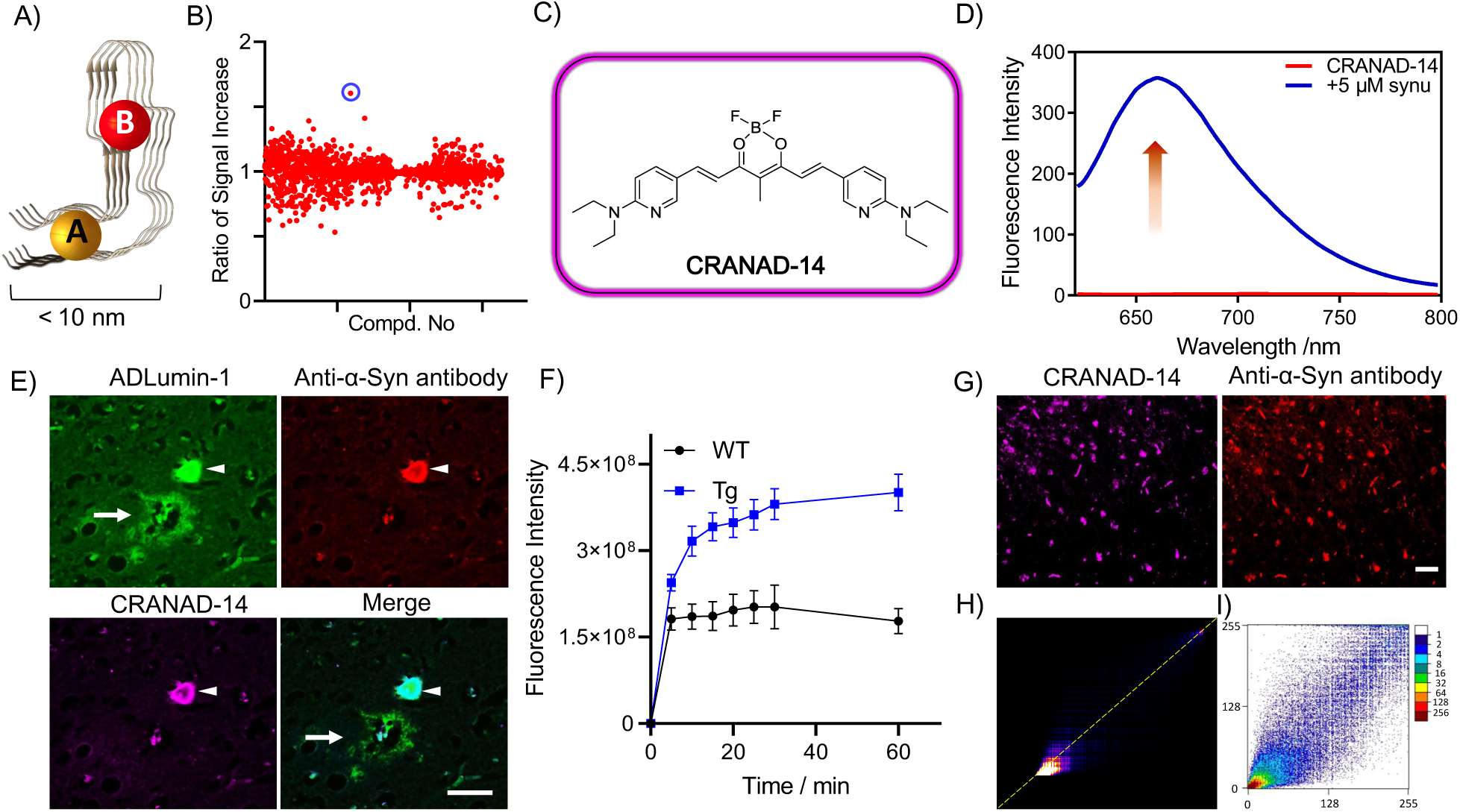
Screening of NIR fluorophores for selective bio-orthogonal-ChRET imaging of misfolded α-syn. (**A**) Diagram of potential probe binding sites (A and B) within α-syn fibrils. (**B**) Fluorescence compound library screening for α-syn aggregates. The ratio of signal increases of fluorescence compounds (5 μM) and 500 nM α-syn aggregates were quantified at different filters (Ex = 605-710 nm, Em = 660-840 nm). CRANAD-14 showed the highest signal increase before and after α-syn aggregates. (**C**) Chemical structure of CRANAD-14. (**D**) Fluorescence spectra of CRANAD-14 (200 nM) and α-syn aggregates (5 μM). The addition of α-syn aggregates showed a 158.14-fold increase in intensity and a 24-nm blueshift of the emission wavelength of CRANAD-14. (**E**) Fluorescence staining of α-syn aggregates in the slides of patients with Parkinson’s disease. CRANAD-14 (magenta color) could stain α-syn aggregates (arrowhead), which were confirmed by subsequently immunostaining with anti-α-syn inclusions antibody (MJFR antibody, red color). CRANAD-14 fails to stain Aβ plaques, which was confirmed by ADLumin-1 staining (green color). (**F**) Time courses of near-infrared fluorescence imaging of 9-month-old A53T transgenic mice and wild-type mice at 0 to 60 min after intravenous injection of CRANAD-14 (4 mg/kg). (**G**) Ex vivo histological observation of A53T transgenic mice (female, 9-month-old, n = 3) after CRANAD-14 injection. A53T mice showed α-syn aggregates labeled by CRANAD-14, which was confirmed by subsequent staining of an antibody that is specific for α-syn aggregates (scale bar = 50 μm). (**H, I**) Colocalization analysis and Pearson’s R value calculation of CRANAD-14 and antibody staining group. Both Colocalization Finder (**H**) and Scatter J (**I**) tests showed good colocalization and the Pearson’s R values for both tests are 0.93, indicating CRANAD-14 could label α-syn aggregates.

However, selective small molecular NIR fluorescence probes for in vivo imaging of α-syn fibrils are scarce (*69, 80*). In this regard, we set out to identify a suitable NIR fluorescence probe that can be used to pair with ADLumin-1 for the biorthogonal ChRET imaging of α-syn. Over the past decades, our group developed a library of various near-infrared (NIR) fluorophores (*38, 39, 42, 81, 82*). To seek α-syn-selective probes, we conducted a plate-based screening against the library by quantifying the fluorescence intensity change before and after adding 50 nM α-syn aggregates (Fig. 6B). Among all the compounds, CRANAD-14 (structure in Fig. 6C, characterization data are shown in Fig. S14-16) showed the most significant fluorescence intensity increase upon interacting with α-syn aggregates. To further validate the binding of CRANAD-14 with α-syn aggregates, the fluorescence spectra of CRANAD-14 were recorded before and after adding the α-syn aggregates. As shown in Fig. S17A, CRANAD-14 alone showed weak fluorescence with an excitation peak at 611 nm, and an emission peak at 685 nm. After binding with α-syn aggregates, the fluorophore shifted its excitation to 604 nm, and emission to 661 nm (Fig. S17B). The chemiluminescence spectra of ADLumin-1 were also recorded with α-syn aggregates. The results indicated good overlap between the chemiluminescence spectra of ADLumin-1 and the excitation spectral of CRANAD-14, and thus enabled ChRET imaging (Fig. S17C). With 5 μM of α-syn aggregates, CRANAD-14 showed a 158.14-fold increase in fluorescence intensity (Fig. 6D). In addition, compared to Aβ aggregates, CRANAD-14 has stronger binding with α-syn aggregates, showing excellent binding affinity (K*d* = 13.03 ± 2.29 nM) (Fig. S18). Furthermore, CRANAD-14 was used to stain brain slides of a PD patient (6-μm-thick, temporal lobe, male, 85 years old), showing excellent labeling of α-syn inclusions and faint staining of Aβ plaques, which were confirmed by subsequent staining of anti-α-syn aggregates antibody and ADLumin-1 (Fig. 6E). In addition, we performed near-infrared fluorescence imaging, and found that CRANAD-14 could differentiate transgenic mice from WT mice, evidenced by significantly higher fluorescence intensity in the brains of the transgenic mice (Fig. 6F). The brains of transgenic and WT mice were subsequently harvested, cut into slides and observed under fluorescence microscopy. To further confirm the substantial labeling of α-syn aggregates, the ex vivo brain tissues were co-stained with an anti-α-syn aggregates antibody. As shown in Fig. 6G, CRANAD-14 and antibody staining group displayed numerous fluorescence labeling of α-syn inclusions and Lewy neurites, while WT mice showed no apparent labeling (Fig. S19). Furthermore, we conducted colocalization analysis of CRANAD-14 and antibody staining group by using Colocalization Finder and Scatter J. As shown in Fig. 6H and I, both tests displayed good colocalization and the Pearson’s R values for both tests are 0.93, indicating CRANAD-14 could label α-syn aggregates. Collectively, these data indicated that CRANAD-14 is specific to α-syn and could be used for bio-orthogonal-ChRET imaging.

### In vitro and in vivo mimic of bioorthogonal-ChRET imaging

To investigate whether in vitro bio-orthogonal-ChRET phenomena could be observed with the ADLumin-1/CRANAD-14 pair, ADLumin-1 in PBS solution was mixed with or without α-syn aggregates in the presence or without CRANAD-14 (Fig. 7A). The samples were put into an IVIS imaging system and scanned at different wavelengths of emission filters (500 nm to 800 nm) with a blocked excitation filter. The data were analyzed by using the spectra unmixing function in the IVIS system. As shown in Fig. 7B, a strong signal turn-on effect with two different spectra was observed in the presence of α-syn aggregates with or without CRANAD-14 (Fig. 7B, C). In contrast, the control groups (no α-syn aggregates) showed very low chemiluminescence signals, which suggests that bioorthogonal-ChRET occurs between ADLumin-1 and CRANAD-14. To further investigate the ChRET phenomenon in a biologically relevant environment, the solution of the ADLumin-1/CRANAD-14 pair was mixed with α-syn aggregates, and the mixture solutions were spiked in mice brain homogenates and imaged with the same parameters. As shown in Fig. 7D and E, the ChRET phenomenon could also be observed in brain homogenates, indicating negligible interference of non-specific interaction to ADLumin-1/CRANAD-14 pair in the biological environment.

**Fig. 7.**
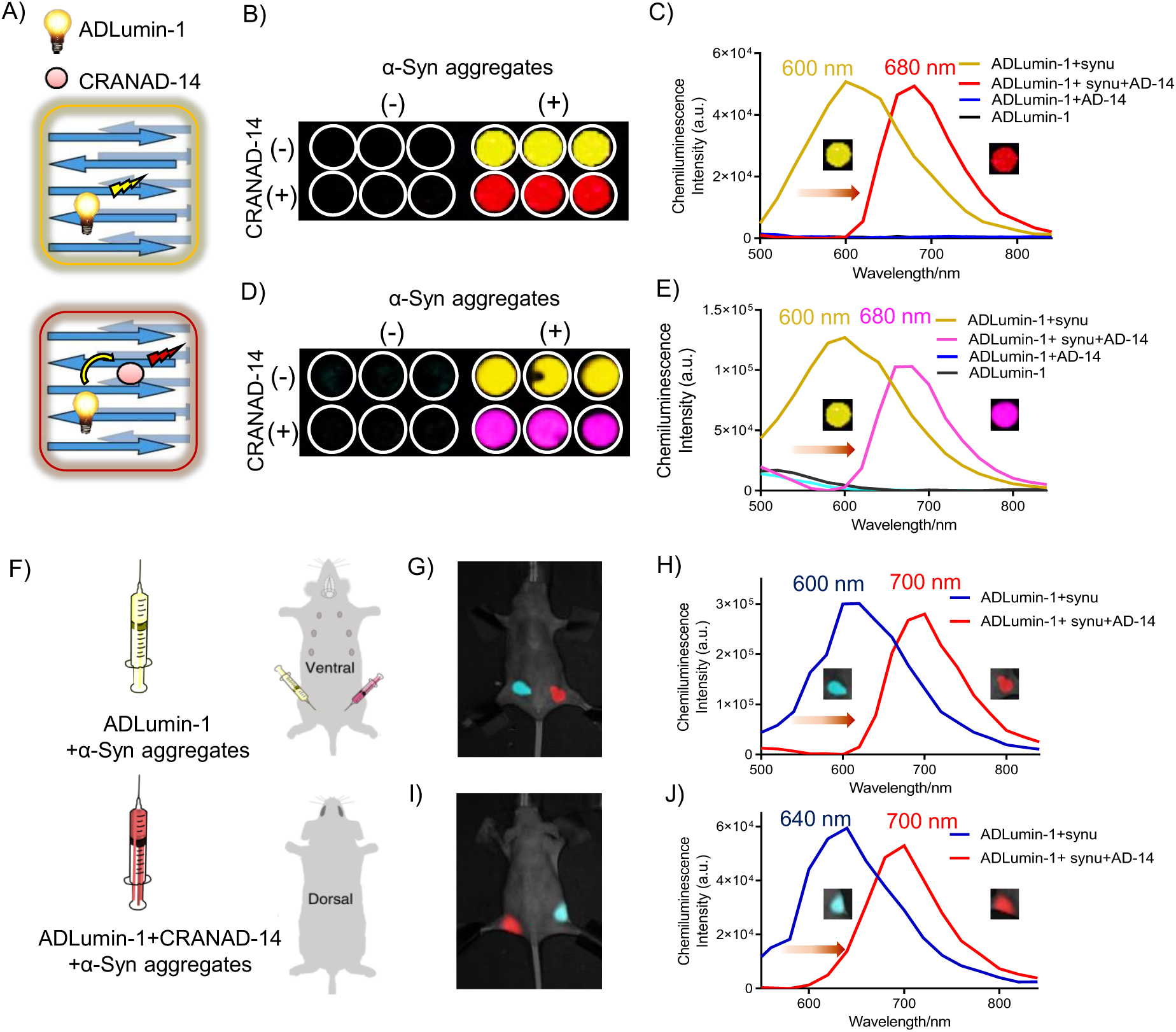
In vitro and in vivo mimic of bioorthogonal-ChRET imaging. (**A**) Diagram of the bio-orthogonal-ChRET strategy. ADLumin-1 can bind with α-syn aggregates and generate chemiluminescence signals. When both ADLumin-1 and CRANAD-14 bind with α-syn aggregates, a ChRET phenomenon will happen, and the CRANAD-14 will be excited and produce signals in long emissions. This process enables selective misfoldon detection based on rigorous proximity requirements (less than 10 nm) and overcomes background interference. (**B**-**E**) Representative spectra unmixing images (**B**, **D**) and spectra of ADLumin-1 (**C**, **E**) with or without α-syn or CRANAD-14 in PBS solution (**B**, **C**) and mouse brain homogenate solution (**D**, **E**). Negligible chemiluminescence signal can be observed without α-syn aggregates. When bound to α-syn aggregates with or without CRANAD-14, ADLumin-1 displayed two emission peaks at 600 nm and 680 nm, respectively. (**F**) Diagram of the in vivo mimic bio-orthogonal-ChRET imaging. ADLumin-1 was mixed with α-syn aggregates with or without CRANAD-14. The mixture was injected to the left and right side of the mice at the ventral hind limb. (**G**-**J**) Representative spectra unmixing images and spectra of injection sites at ventral (**G**, **H**) and dorsal (**I**, **J**) positions. When bound to α-syn aggregates with or without CRANAD-14, ADLumin-1 displayed two emission peaks at 600 nm and 700 nm for ventral position, and 640 nm and 700 nm for dorsal position, respectively.

The significant bio-orthogonal-ChRET phenomenon of the ADLumin-1/CRANAD-14 pair with α-syn aggregates inspired us to investigate in vivo. A mimic study was first conducted in nude mice. The solution of ADLumin-1 and α-syn aggregates with or without CRANAD-14 were subcutaneously injected into different sites of nude mice at ventral position (Fig. 7F), and the mice were imaged under the IVIS system with a blocked excitation filter and emission filters (500 nm to 800 nm). As shown in Fig.7 G-J, two distinct emission spectra could be observed, with peaks at 600 nm (without CRANAD-14) and 700 nm (CRANAD-14) when the mouse was imaged from the ventral position. Interestingly, if the same mouse were imaged from the dorsal side, the spectral peaks were red-shifted to 640 nm (without CRANAD-14) and 700 nm (with CRANAD-14), respectively. The red-shifting is likely due to the absorption of short emission wavelength in biological tissue that displayed pseudo peaks as longer emission wavelength. Taken together, our data confirmed that the bio-orthogonal-ChRET could be observed in biologically relevant environments.

### In vivo bio-orthogonal-ChRET Imaging with ADLumin-1/CRANAD-14 pair

To validate the feasibility of in vivo ChRET imaging, A53T transgenic mice and WT control mice were intravenously injected with CRANAD-14 (4 mg/kg) and intraperitoneally injected with ADLumin-1 (8 mg/kg) at 30 min postinjection of CRANAD-14. The chemiluminescence signals of mice were scanned from 500 nm to 800 nm. After the completion of imaging acquisition, automatic spectral unmixing was applied (Fig. 8A). The resulting unmixing images and quantitative chemiluminescence signal showed higher chemiluminescence intensity from the transgenic mice brains compared to the WT mice (Fig. 8B, C). Furthermore, the highest ratio of transgenic mice to WT mice was quantified. The bio-orthogonal-ChRET strategy displayed 2.98 ± 0.27-fold higher signals in the A53T mice group than WT mice. This difference is larger compared to either fluorescence imaging modality by using CRANAD-14 (2.26 ± 0.25) or chemiluminescence imaging modality by using ADLumin-1 (2.06 ± 0.57) (Fig. 8D). Furthermore, to validate that the ADLumin-1/CRANAD-14 pair is selective for detecting α-syn in vivo, we used 5xFAD mice as the control group and imaged with the same protocol. The normalized in vivo spectra (chemiluminescence intensity at different emission wavelengths) of the 5xFAD mice brains are different from A53T mice (Fig. 8E). After subtracted with the in vivo spectra of WT mice, 5xFAD mice showed peaks at 600 nm while A53T mice showed peaks at 700 nm (Fig. 8F). As 700 nm is the wavelength after bio-orthogonal-ChRET in the presence of α-syn fibrils (Fig. 7), this indicated that bio-orthogonal-ChRET occurred only in the presence of misfolded α-syn in the A53T mice brain that generated the redshift of emission wavelength but not in the 5xFAD mice brains that contain misfolded Aβ.

**Fig. 8.**
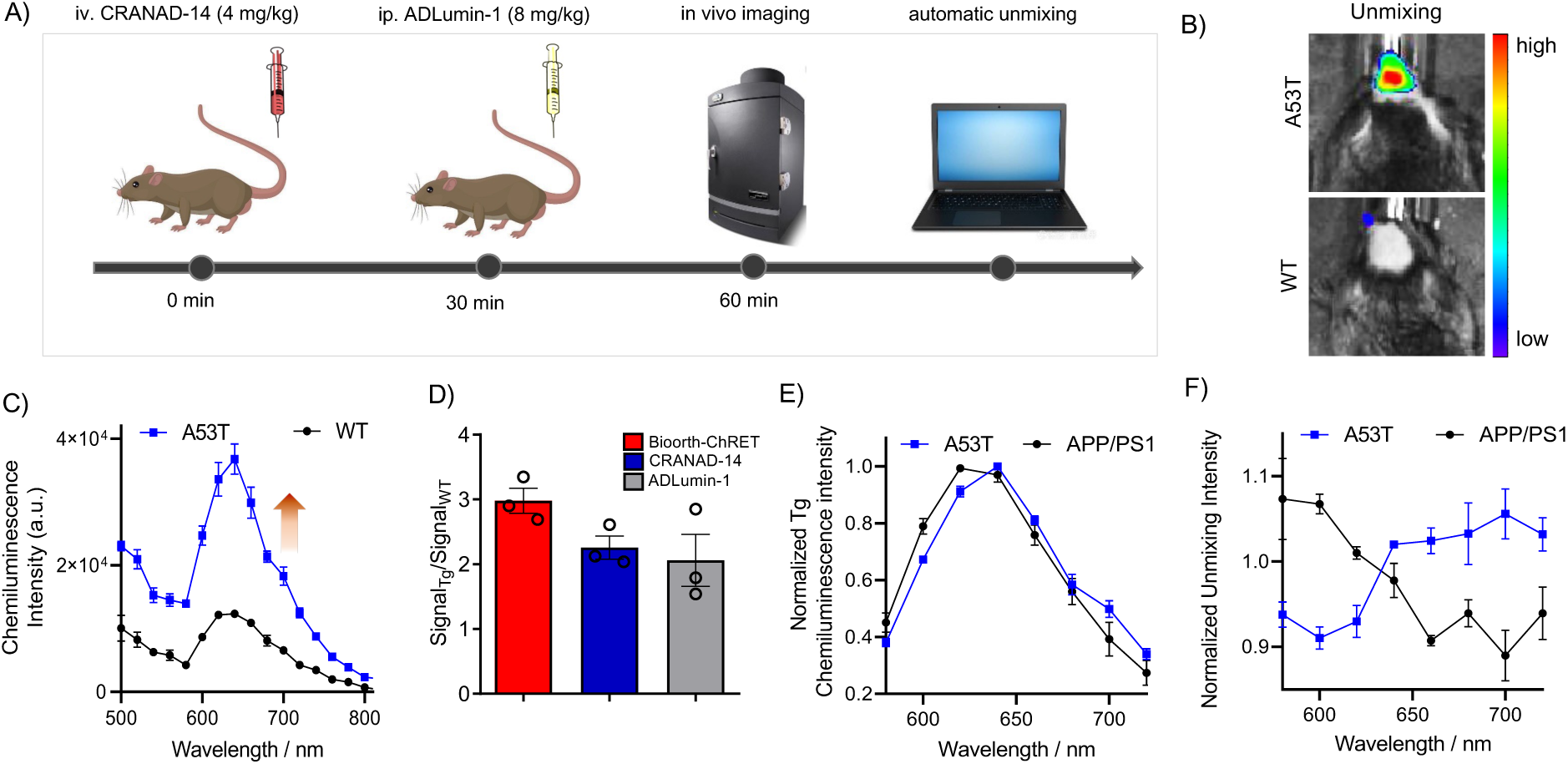
In vivo bio-orthogonal-ChRET Imaging with ADLumin-1/CRANAD-14 pair. (**A**) Workflow of in vivo bio-orthogonal-ChRET imaging. A53T transgenic mice (female, 9-month-old, n = 3) and age-matched wild-type control mice (female, 9-month-old, n = 3) were intravenously injected with CRANAD-14 (4 mg/kg). After 30 min post-injection, the mice were intraperitoneally injected with ADLumin-1 (8 mg/kg). The mice were put into the IVIS imaging system, and the chemiluminescence signals from 500 nm to 800 nm were quantified, automatically unmixed, and analyzed. (**B**) Representative images of transgenic and wild-type mice after automatically unmixing. (**C**) Quantitative chemiluminescence signal of transgenic mice (Tg) and wild-type mice (WT) at different emission wavelength filters. (**D**) Comparison of the signal ratio of transgenic mice and wild-type mice by using the bio-orthogonal-ChRET, ADLumin-1, or CRANAD-14. **E**) Normalized in vivo spectra (chemiluminescence intensity at different emission wavelengths) of the 5xFAD and A53T mice brains. **F**) Normalized in vivo spectra of 5xFAD and A53T mice after subtracted with the in vivo spectra of WT mice. 5xFAD mice showed peaks at 600 nm while A53T mice showed peaks at 700 nm, indicating that bio-orthogonal-ChRET occurred only in the presence of misfolded α-syn in the A53T mice brain.

Collectively, these data support the potential of ADLumin-1 based bio-orthogonal-ChRET strategy for in vivo imaging of cerebral α-syn fibrils with improved selectivity and signal increase.

## Discussion

Aggregating of misfolded β-sheet is a general process for various chronic diseases (*83*). However, general methods for detecting such aggregation are limited, particularly for in vivo studies. ThT fluorescence is the most used method for quantifying misfolded β-sheets, but it is limited to in vitro tests, such as pure solutions, biofluids, tissue homogenates, and tissue staining (*29*). CD and FT-IR are often used for identifying β-sheet conformation but are restricted to pure protein structure studies. Immunoassay is also the most commonly used approach for detecting specific β-sheets; however, designated antibodies are needed (*84, 85*). Clearly, a generic probe that can detect various misfolded proteins in vivo is highly desirable.

With the advance of optical imaging, numerous fluorescence probes have been developed for detecting various misfoldons, including Aβ and tau (*38, 40, 86–88*). However, chemiluminescence probes are rare for this purpose (*42, 71*). In this report, we demonstrated a new approach to monitoring proteinopathies with our chemiluminescence probe ADLumin-1. To validate that ADLumin-1 can be considered a generic probe for misfoldons, we tested the probe with 7 representative misfoldons in vitro and transgenic mice models containing misfolded tau, TDP-43 or α-syn, which are the key biomarkers of AD, ALS, and PD. Indeed, our data suggested that ADLumin-1 was capable of in vivo imaging of these misfoldons.

Compared to the gold standard dye ThT and previous imaging probes, ADLumin-1 has several advantages: 1) It has high sensitivity, due to its chemiluminescence features, such as no requirement of excitation light, and minimal interference from biological autofluorescence (*89, 90*). 2) Our in vitro studies suggested that ADLumin-1 could be sensitive towards oligomers, whereas ThT is known to be insensitive to oligomers. 3) ADLumin-1 has the capacity for in vivo imaging applications. In contrast, ThT’s poor brain permeability, short emission wavelength, and low SNR limit its in vivo applications (*29*). 4) The application of ADLumin-1 is simple, fast, and versatile. ADLumin-1 can be applied for various proteinopathy animal models via intravenous and intraperitoneal injection. In contrast, most fluorescence probes can only be administrated intravenously to animals (*89*).

The interacting mechanism of ThT for various β-sheets has triggered increasing interest, but there is no clear answer yet (*91*). For ADLumin-1, our molecular docking results indicated the interaction of ADLumin-1 to the long-axis of the misfolded β-sheet structure. The results suggested that its detecting capacity originated from its binding to hydrophobic pockets that are formed by β-sheets, particularly by Ala, Val, Ile, Leu, and Phe, which are the driving amino acids to form misfolded β-sheets (*26, 27*). In addition, ADLumin-1 showed distinct signal decay half-life time when bound with misfoldons, and this feature could be used for misfoldon differentiation. However, we didn’t intend to explore the detailed mechanism in this study.

With α-syn as an example, we further demonstrated both the generic and selective detection capability of ADLumin-1. Recently, based on the fluorescence signal of ThT, PMCA and RT-QuIC were developed to report the concentrations of prion, Aβ, and other misfoldons in human biofluids via cyclic signal amplification (*33–35*). These methods showed ultra-sensitive detecting capability and are promising to translate to clinical diagnosis (*37, 92–94*). Interestingly and importantly, we also demonstrated that ADLumin-1 could be used as a highly sensitive and selective detecting probe for PMCA with α-syn, and the sensitivity of ADLumin-1 was higher than that of ThT. Our preliminary in vitro results suggested that ADLumin-1 could be an excellent alternative for ThT in biofluid detection.

To visualize α-syn pathology in vivo, we first revealed that ADLumin-1 could be used for visualizing the seeding of α-syn in the brain via 3D reconstruction. Given that the propagation of α-syn in brains plays an important role in synucleinopathy, it is highly desirable to monitor the propagation longitudinally in vivo. Based on our results, ADLumin-1 has the potential to monitor cerebral α-syn misfolding and propagation via 3D tomography imaging (*46, 95*). For A53T transgenic mice, we monitored the α-syn progression in mice from 4-month-old to 12-month-old of ages. ADLumin-1 showed increasing signals in transgenic mice compared to WT mice. Interestingly, previous studies indicated the appearance of α-syn inclusions in the brain of 9-month-old A53T mice, and more recent evidence indicated that α-syn pathology occurs even as early as the 4-month age of A53T mice (*96*). Our imaging results may indicate protein misfolding behaviors at the early stage of A53T mice, and further investigations are needed.

Non-conjugated bio-orthogonal-ChRET imaging is a new concept that we recently reported (*42, 79, 97*). This study is the first example of bioorthogonal-ChRET application for α-syn aggregates. The ChRET imaging utilized a nonconjugated ChRET pair of a chemiluminescence probe and a NIR fluorescence probe. As the energy transferring requires a match of emission/excitation wavelength and the proximity of two probes, undesirable signals caused by non-specific binding can be considerably reduced through this process. We envisioned that utilizing our probe with bioorthogonal-ChRET to differentiate distinct conformers would be highly desirable in future research. This is due to the heterogeneity of misfolded conformers occurring in various neurodegenerative diseases or progression stages, and thus the development of imaging technologies that can selectively detect these distinct conformers in vivo is crucial and in high demand (*56, 98–100*). However, developing a highly selective probe is challenging due to the structural similarity of misfolded protein and the rigorous requirements of brain penetration (*69, 80*). Thus, the probe development normally takes years, and few could be widely used in preclinical and clinical research. Instead, our technology provides an alternative strategy by using proximity chemistry. With the advance of imaging probes, spectra unmixing algorithm, and 3D reconstruction technology, the principle of bio-orthogonal-ChRET could be used for further differentiation of broad spectra of proteinopathies in vivo.

Despite the diagnostic value, we also envisioned that the binding of ADLumin-1 may modulate misfoldon functions by disrupting β-sheet formation and has therapeutic potential for proteinopathies. Further study is undergoing in our group.

In summary, we first used model peptides to validate that ADLumin-1 has high specificity for β-sheet fibrils and showed up to 127.73-fold higher SNR compared to gold standard dye ThT. Next, we demonstrated the generic detecting capability of ADLumin-1 for various misfoldons with excellent signal amplification. We also proved the in vivo imaging capability of ADLumin-1 to visualize α-syn, tau, and TDP-43 pathology in trangenic mice models. By using α-syn as a model biomarker, we demonstrated the selective detecting capability of ADLumin-1 by combining PMCA technology in vitro and bio-orthogonal-ChRET imaging strategy in vivo. Given that our detection procedure is very sensitive, versatile, fast, and low cost, we believe that ADLumin-1 is an important complementary tool for detecting misfoldons both in vitro and in vivo. ADLumin-1 has the potential to be widely applied in pre-clinical and clinical detections of crucial biomarkers of the most influential neurodegenerative diseases, including AD, PD, and ALS, and facilitate disease early detection, etiology elucidation, and drug discovery.

## Materials and Methods

### General Information

All reagents were commercial products and used without further purification. Palmitic peptide PA-K (sequence: KKK), PA-K2 (sequence: VVVAAAKKK), PA-E (sequence: EEE), PA-E2 (sequence: VVVAAAEEE), CATCH+ (QQKFKFKFKQQ) and CATCH-(EQEFEFEFEQE) were purchased from Genscript. Synthetic α-syn and Aβ_1-40_ peptide were purchased from rPeptide. Tau preformed fibrils were purchased from StressMarq. Transthyretin protein was purchased from Sinobiological. Prion protein and amylin were purchased from Sigma-Aldrich. Anti-Alpha-synuclein aggregate antibody [MJFR-14-6-4-2] (ab209538), Anti-Tau (phospho S404) antibody [EPR2605], (ab92676), Anti-TDP43 antibody [EPR5810] (ab109535) were purchased from Abcam. Alexa Fluo555-conjugated goat anti-rabbit IgG (H+L) secondary antibody was purchased from Thermofisher (A-21428). Cerebrospinal fluid (CSF) and paraffin brain tissue samples are commercially available products from Innovative Research or Novus Biologicals. B6;C3-Tg(Prnp-SNCA*A53T)83Vle/J mice (RRID:IMSR_JAX:004479), STOCK Mapttm1.1Cole/J (RRID:IMSR_JAX:033373), B6.Cg-Tg(Prnp-TARDBP*A315T)95Balo/J (RRID:IMSR_JAX:010700), and wild-type mice were purchased from the Jackson lab. All animal experiments were approved by the Institutional Animal Use and Care Committee (IACUC) at Massachusetts General Hospital (approval number: 2011N000161), and carried out in accordance with the approved guidelines.

### Protein Aggregates Preparation

α-syn protein (70 μM) was dissolved in PBS buffer (1X, pH = 7.4) and incubated for 72 h with constant shaking. Other misfoldons were prepared by following a similar incubation protocol as α-syn protein, and the successful formation of protein aggregates was confirmed by the standard dye of thioflavin T (15 μM). The samples are characterized by following previously reported methods (*56, 63, 101*). The detailed preparation methods are in SI.

### Transmission Electronic Microscopy (TEM) Observation

Palmitic peptide samples were dissolved in PBS buffer (1X, pH = 7.4) and incubated for 72 h with constant shaking. CATCH+/− pairs were mixed before TEM observation. A drop of the sample was placed onto a TEM mesh grid, and a negative staining reagent was applied. The morphology images were captured by a JEOL JEM-1011 TEM instrument.

### Circular Dichroism (CD) Spectra Analysis

Palmitic peptide samples were diluted with 10 mM phosphate buffer to give a final concentration of 10 μM. Samples were pipetted into a 10 mm patch cuvette, and the CD spectra were recorded with a Jasco J-815 Spectropolarimeter. The bandwidth was 1 nm, and the scanning started at 260 nm and ended at 190 nm. Data pitch was 0.1-0.5 nm. The scanning speed was 50-100 nm/min. Each sample was accumulated 5 times, and the signals were averaged to give final spectra.

### Chemiluminescence Detection with ADLumin-1

Protein samples were dissolved in PBS buffer (1X, pH = 7.4). To a solution of protein aggregates, an ADLumin-1 DMSO stock solution was added (final concentration: 15 μM in 5% DMSO/PBS buffer). The mixture was immediately transferred to a black 96 or 384 well and observed with an IVIS®Spectrum imaging system (Caliper Life-Sciences, Perkin Elmer) with an open filter. The signals were recorded from 5 min to 200 min, and the region-of-interests (ROI) were quantified by LivingImage® 4.2.1 software. The signal increase was calculated by the ratio of the highest chemiluminescence intensity of samples to the blank control group. Half-life times of the signal decay were calculated by using GraphPad Prism 8.0 with nonlinear regression decay.

### Molecular Docking

The ligand structure was built and optimized using IQmol (IQmol, version 2.15.3). The receptor structures (PDB: 6LNI, 6VW2, 5O3L, 6PEO, 5OQV, 7OB4) were then loaded in UCSF Chimera (UCSF Chimera, version 1.15). The receptor structures were then prepared for docking through Dock Prep, which added hydrogen atoms and assigned atomic partial charges. For standard residues, AMBER ff14SB charges were used, while Gasteiger charges were assigned for nonstandard residues. AutoDock Vina 1.2 was used to dock the ligand to the receptor using the default settings. The initial docking was conducted by constructing a box encompassing the entire protein and finding the best (global) docking position within that box. Ligand plots were generated using Schrödinger Maestro (Schrödinger Maestro, version 13.0.137) by using a ligand-protein complex of each of the best docking results for each protein that was made using Chimera.

### Binding Affinity

Different concentrations of ADLumin-1 (0-40 μM, 10% DMSO/PBS) were mixed with α-syn fibrils (250 nM) or PBS solution. The mixture was immediately transferred to a 96-well black plate, and the chemiluminescence intensity was recorded using an IVIS imaging system. Dissociation constants (Kd) were calculated using GraphPad Prism 8.0 with nonlinear one-site binding regression. The binding affinity of CRANAD-14 for α-syn aggregates or Aβ aggregates was measured by the following procedure: Aβ aggregates (500 nM), α-syn aggregates (500 nM) or PBS solution were mixed with different concentrations of CRANAD-14 (0-500 nM). The mixture was added into a quartz cuvette, and the fluorescence intensities were quantified by an F-7100 fluorescence spectrometer (Hitachi, Japan). Dissociation constants (Kd) were calculated using GraphPad Prism 8.0 with nonlinear one-site binding regression.

### Protein Misfolding Cyclic Amplification (PMCA) Assay

Seed samples were prepared by spiking α-syn aggregates (70 fM, final concentration, sonicated before adding) into cerebrospinal fluid (CSF). 40 μL of seed samples or CSF were added to 200 μL1mg/ml α-syn monomer solution in 100 mM PIPES, pH 6.5, and 500 mM NaCl. Samples were subjected to cyclic amplification (1 min shaking followed by 29 min incubation) at 37 °C (*37*). After cyclic amplification, samples were transferred to a 384-well black plate, followed by adding ADLumin-1 (15 μM in 5% DMSO/PBS buffer) and measured with an IVIS®Spectrum imaging system (Caliper Life-Sciences, Perkin Elmer) equipped with a bioluminescence filter. The signals were recorded at 5 min to 200 min and the ROIs were quantified by LivingImage® 4.2.1 software.

### Histological Staining

Paraffin brain tissue sections from a Parkinson’s disease (PD) patient were obtained from Novus Biologicals (6-μm-thick, temporal lobe, male, 85 years old) and were deparaffinized in xylene, 100% ethanol, 95% ethanol, and dd-water. Brain sections were incubated with 5 μM of the probe for 30 min at room temperature, and were then rinsed with 40% ethanol (2 × 1 min) and H_2_O (2 × 30 s). Fluorescence images of the sections were obtained under a Nikon Eclipse 50i microscope. The presence of α-syn was confirmed by subsequent immunostaining of the same section with recombinant anti-α-syn antibody [MJFR] (Abcam) and Alexa Fluo555 labeled secondary antibody.

### 3D Imaging of Cerebral α-syn

Stereotactic injection surgery was conducted by using C57BL/6J mice (female, 9-month-old, n = 5). Freshly prepared α-syn aggregates (62.5 μM, 400 nL, 0.02 nL/sec) were injected into the substantial nigra region of mice (AP: −3.29 mm; ML: ±1.25 mm; DV: −4.35 mm). After recovering for 1 month, the mice were anesthetized and head-shaved, followed by scanning background signals with an IVIS®Spectrum animal imaging system (Caliper Life-Sciences, Perkin Elmer). A solution of ADLumin-1 (8 mg/kg) was freshly prepared in the mixed solvent (15% DMSO, 15% cremorphor and 70% PBS). 100 μL of ADLumin-1 was intravenously injected to the mice, followed by acquiring CT images. At the same time, the chemiluminescence signals were recorded with the blocked excitation and several filters of emission (560 nm to 660 nm) under the DLIT mode of the IVIS system. The images were automatically reconstructed by a LivingImage software® 4.2.1.

### In Vivo Imaging

Female transgenic α-syn A53T mice (Tg) and age-matched wild-type mice (WT) (n = 5, 4-12 months old) were anesthetized and shaved in the brain region. Background signals were recorded by using an IVIS®Spectrum animal imaging system (Caliper Life-Sciences, Perkin Elmer). 100 μL freshly prepared ADLumin-1 (8 mg/kg, 15% DMSO, 15% cremorphor, and 70% PBS) was intraperitoneally injected into the mice, and chemiluminescence signals were recorded with the parameter of blocked excitation and open filter of emission by using IVIS system. Same imaging protocol was used for P301L tau (n = 3, 10 months old) and A315T transgenic mice (n = 3, 6 months old). For CRANAD-14 imaging, 4 mg/kg CRANAD-14 in a mixed solvent (15% DMSO, 15% cremorphor, and 70% PBS) was intravenously injected to 9-month-old A53T or wild-type mice. Then, the mice images were acquired from 0 to 60 min under an excitation wavelength of 605 nm and emission wavelength of 680 nm. Data analysis was conducted by calculating the average radiance [p/s/cm^2^/sr] of ROI with LivingImage® 4.2.1 software. The signal intensity at each time point was normalized by deducting the background signal.

### Fluorophore Screening

CRANAD-X library with 137 small molecules were synthesized by our lab. 50 μL of α-syn aggregates (50 nM) or PBS solution were added to each well of a black 96-well plate, followed by adding a 2.5 μL solution of CRANAD-X (final concentration: 5 μM, in 5% DMSO/PBS). After incubating for 30 min in a dark room, the plate with or without α-syn aggregates were scanned by an IVIS®Spectrum imaging system (Caliper Life-Sciences, Perkin Elmer) equipped with fluorescence filters (excitation = 605∼710 nm; emission = 660∼840 nm). The signal was quantified by the LivingImage software, and the fluorescence intensity of test groups at each filter was normalized by deducting the signal of blank control groups.

### Bio-orthogonal-ChRET

For in vitro studies, 200 μL 25uM α-syn aggregates or PBS solution was mixed with 5 μL CRANAD-14 (final concentration: 25 uM in 2.5% DMSO/PBS) or DMSO solution. The samples were incubated for 30 min, followed by the addition of ADLumin-1 (15 μM in 5% DMSO/PBS buffer). For in vivo mimic studies, 45 μL 25uM α-syn aggregates were added into 2.5 μL 15 μM ADLumin-1 solution and mixed with or without 2.5 μL 25 uM CRANAD-14 solution. Nude mice (female, 6 weeks old, n = 3) were anesthetized, and the left and right hind limbs were subcutaneously injected with different sample solutions. For in vivo studies, head-shaved transgenic A53T and wild-type mice (female, 9-month-old, n = 3) were intravenously injected with CRANAD-14 (4 mg/kg). After 30 min, the mice were intraperitoneally injected with 8 mg/kg ADLumin-1. The samples or mice were placed into an IVIS system, and the sequence images were acquired with open filter and different emission filters (open filter and 500∼800 nm). ROI was drawn around the wells or mice brains for quantifying the signal. The data were analyzed by Living Image 4.2.1 software with guided spectra unmixing method and default parameters.

### Statistical analysis

Quantitative data shown as mean ± s.e.m. were analyzed by GraphPad Prism 8.0 software. P values were determined by unpaired two-tailed Student’s t-tests. The differences were considered significant when P ≤ 0.05.

## Funding

National Institutes of Health grant R01AG055413 (NIH, CR)

National Institutes of Health grant R01AG083759 (NIH, CR)

National Institutes of Health grant R01AG085562 (NIH, CR)

National Institutes of Health grant R21AG059134 (NIH, CR)

National Institutes of Health grant R56AG059814 (NIH, CR)

National Institutes of Health grant R21AG078749 (NIH, CR)

National Institutes of Health grant S10OD028609 (NIH, CR)

Natural Science Foundation of Chongqing CSTB2024NSCQ-MSX0365 (NSCQ, BZ)

Chongqing Science and Health Joint Medical Research Project 2025MSXM061 (BZ)

## Author contributions

Writing—original draft: CR, BZ

Conceptualization: CR, BZ, CZ

Investigation: CR, BZ, AY, RV, JW, EL, FY

Writing—review & editing: CR, BZ, RV, SK, FL, JZ, JY, RT, YJ, EL, HW, FY

Methodology: CR, BZ, RV, SK, HH, JW, EL

Resources: CR, BZ, CZ, XM, SK, FL, YJ, RT, YS

Funding acquisition: CR, BZ

Data curation: CR, BZ, AY, HW

Validation: CR, BZ, HH, HW

Supervision: CR, BZ

Formal analysis: CR, BZ, AY, CZ, RV, YJ

Project administration: CR, BZ, CZ, JZ, JY, HW

Visualization: CR, BZ, RV, SK, YJ

## Competing interests

All authors declare they have no competing interests.

## Data and materials availability

All data are available in the main text or the supplementary materials.

## Notes

### Competing Interest Statement

The authors have declared no competing interest.

