## Supplemental Materials for "Detection of misfolded proteins in neurodegenerative disease models with a highly sensitive chemiluminescence probe"

Biyue Zhu *et al.*

**This PDF file includes:**

Supplementary Text
Figs. S1 to S19
Tables S1 to S3

### Supplementary Text

#### Protein Concentration Quantification.

The protein concentrations were quantified by following the standard protocols of Pierce™ BCA Protein Assay kit. Briefly, triplicates of 25 μL bovine serum albumin solution (20–2,000 μg/mL) or protein samples were added into a 96-well plate, followed by adding 200 μL BCA working reagent and shaking for 30 s. The plate was covered and incubated at 37 °C for 30 min. The absorption was measured by using a microplate reader at 562 nm. The blank standard replicates absorbance at 562 nm was subtracted, followed by plotting the standard curve with BSA standard absorbance intensities and known concentrations. The unknown sample concentrations were calculated based on the standard curve.

#### Protein Fibrils Preparation.

All protein samples are in wild-type form. For ThT comparison test, tau preformed fibrils were purchased from StressMarq (SPR-480). Recombinant α-synuclein (α-syn, rPeptide, S-1001-2), Aβ (rPeptide, A-1001-1), or TDP-43 (Abcam, ab224788) proteins were dissolved in PBS buffer (1X, pH 7.4), followed by filtered through 0.2 μm filters (Sigma-Aldrich). The filtrate was transferred into 0.5 mL EP tubes and incubated for 72 h with 1000 rpm constant shaking. Samples were centrifuged at 20,000 g, 4 °C for 10 min to separate the soluble species and collected insoluble fibrils. The successful formation of protein fibrils was confirmed by TEM and the standard dye of thioflavin T (ThT).

#### Fibril Polymorphs Preparation.

Different α-syn fibril polymorphs were prepared in buffers with (S fibrils) or without (NS fibrils) sodium chloride. For S fibrils, 69 μM α-syn proteins were dissolved in 20 mM Tris-HCl and 100 mM NaCl (pH 7.4). For NS fibrils, α-syn proteins were dissolved in 20 mM Tris-HCl (pH 7.4). Both sample solutions were incubated at 37 °C with continuous shaking at 600 rpm for 7 days (*56*). Different Aβ fibril polymorphs were prepared in buffers with (S fibrils) or without (NS fibrils) sodium chloride. For S or NS fibrils, 100 μM Aβ proteins were dissolved in 50 mM sodium borate (pH 9.0), with (S) or without (NS) 500 mM sodium chloride. Both sample solutions were incubated at 37 °C for 14 days (*54*). Different tau fibril polymorphs (SPR-329, H fibrils; SPR-463, NH fibrils) were purchased from Stressmarq. Different TDP-43 fibril polymorphs were prepared by dissolving 20 μM TDP-43 proteins in 10 mM HEPES with (S) or without (NS) 100 mM sodium chloride (pH 7.4) and incubated for 5 days with continuous shaking at 1000 rpm (*57*).

The samples are characterized by following previously reported methods (*56, 63, 101*). After preparation, all samples were centrifuged at 20000 g for 10 min at 4 °C to separate the soluble species and insoluble fibrils. The protein concentrations were quantified by following the standard protocols of Pierce™ BCA Protein Assay kit. The morphology of each fibril sample was observed by TEM/AFM imaging.

#### Oligomers Preparation.

Unmodified human recombinant α-syn oligomers (SPR-484), dopamine-induced human recombinant α-syn oligomers (SPR-466), human synthetic amyloid beta 1-42 oligomers (SPR-488), and human recombinant tau-441 (2N4R) wild-type oligomers (SPR-497) were purchased from StressMarq. The protein concentrations were confirmed by following the standard protocols of Pierce™ BCA Protein Assay kit. The quality control of oligomers were confirmed by TEM and size exclusion chromatography (SEC).

#### TEM Analysis.

5 μL protein sample solution was dropped onto a mesh-containing copper grid and air-dried. The grid was washed with water, followed by negative staining with 1% phosphotungstic acid (PTA)

or 2% uranyl acetate solution. The samples were observed by JEOL JEM-1011 instrument and TEM images were captured.

**Molecular Docking with Different Protein Fibril Polymorphs or Strains.**

Different protein fibril polymorphs or strains were extracted from PDB databank, including human-derived α-syn (PDB: 6XYO, 6XYQ, 8A9L), recombinant α-syn (PDB: 6A6B, 6SSX, 6PEO), human-derived Aβ (PDB: 8QN6, 8QN7, 2M4J), recombinant/synthetic Aβ (PDB: 2NAO, 5OQV, 2LMO), human-derived tau (PDB: 6NWP, 6NWQ, 5O3L), recombinant tau (PDB: 6QJH, 6QJM, 6QJQ), human-derived TDP-43 (PDB: 8CG3, 8CGG, 7PY2), recombinant TDP-43 (PDB: 8QX9, 8QXA, 8QXB). ADLumin-1's structure was built and optimized using IQmol (IQmol, version 2.15.3). The protein structures were then loaded in UCSF Chimera (UCSF Chimera, version 1.15) and were then prepared for docking through Dock Prep, which added hydrogen atoms and assigned atomic partial charges. For standard residues, AMBER ff14SB charges were used, while Gasteiger charges were assigned for nonstandard residues. AutoDock Vina 1.2 was used to dock the ligand to the receptor using the default settings. The initial docking was conducted by constructing a box encompassing the entire protein and finding the best (global) docking position within that box. Ligand plots were generated using Schrödinger Maestro (Schrödinger Maestro, version 13.0.137) by using a ligand-protein complex of each of the best docking results for each protein that was made using Chimera.

**Sedimentation Assay.**

The sedimentation assay was conducted by following previously reported protocols (*74*). 70 μM α-syn samples were dissolved in PBS buffer (pH = 7.4) and filtered through a 100 KDa MW filter. 4 mM Apigenin or Resveratrol were dissolved in DMSO. 245 μL α-syn samples were mixed with 5 μL Apigenin, Resveratrol or DMSO and shaken for 0, 12, 24, 48, 72 h at 1000 rpm on an orbital shaker. The samples at the final time point (72 h) were directly subjected to TEM imaging to observe the protein morphologies. The samples at each time point were collected and subjected to centrifuge at 20,000 g, 4 °C for 10 min. After carefully separating the soluble and insoluble species, the soluble species (supernatant) were subjected to gel electrophoresis to investigate the remaining soluble fractions, while the insoluble species (pellets) were used for ADLumin-1 detection. For gel electrophoresis, the aliquots of soluble species at each time point were run on 15% SDS-PAGE gel by following the manufacturer's protocols, and the visualization of soluble bands was conducted by subsequent silver staining with Pierce™ silver stain kit. The remaining fractions of soluble protein were determined by quantifying the grey value of each band with Image J software.

**Colocalization Analysis.**

Pathological images captured at different fluorescence channels were loaded into Image J software. The image channels were split, and the images of probe staining and antibody staining were analyzed by using Coloc 2, colocalization and Scatter J plugins with default parameters. The Pearson's values were calculated, and the scatter plots were exported.

**Seeding and Cyclic Amplification Assay.**

As chemiluminescence is different from fluorescence, the form is not reversible upon emitting light, while fluorescence is reversible and can be repeatedly excited, thus a modified seeding and cyclic amplification assay was conducted to compare the detection capability of ThT and ADLumin-1. α-syn protein was dissolved in PBS buffer (pH = 7.4) with 170 mM NaCl and 0.0015% SDS and passed through filters with 100 kDa cut-off to remove any spontaneously formed aggregates. The α-syn protein concentration was quantified by BCA assay before the test. 85 μL α-syn solution (70 μM, final concentration) was added into 5 μL healthy control CSF with or without α-syn seeds (35 pM, final concentration). The samples were added to a 96-well black plate with transparent bottom and separated into ThT group and ADLumin-1. 10 μL ThT (15 μM, final concentration) was added into each well of ThT group before the cyclic amplification. Blank control group was also prepared. The plate was sealed with clear film and subjected to a Synergy H1 microplate reader and conducted cyclic amplification at 42 °C (548 cpm shaking for 1 min, 14 min incubation) for 80 h. The fluorescence intensity of ThT was detected from the bottom of the plate by setting fluorescence parameters with excitation at 440 nm and emission at 485 nm. After cyclic amplification, 10 μL freshly prepared ADLumin-1 in DMSO solution (15 μM, final concentration) was added into ADLumin-1 group and immediately detected by a microplate reader with chemiluminescence detecting mode with default parameters from 0 to 180 min. The fluorescence or chemiluminescence intensity was plotted by using GraphPad software.

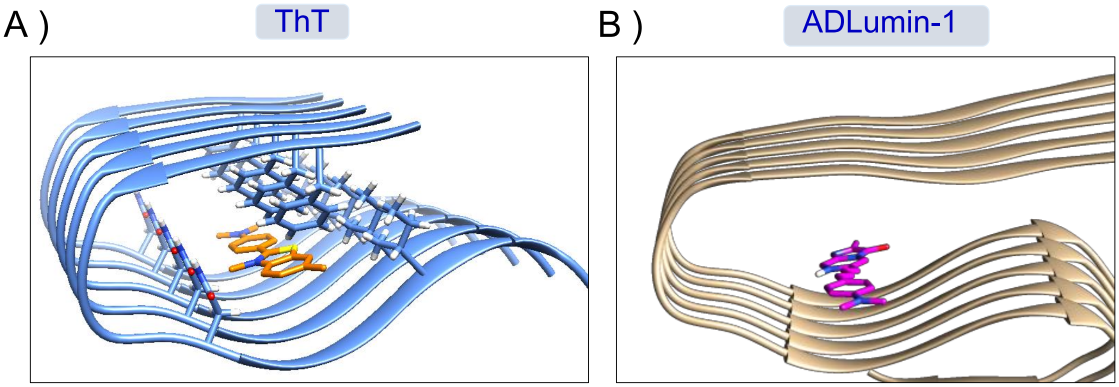


**Fig. S1.** Binding pose of ThT and ADLumin-1 to Aβ fibrils (PDB: 5OQV). Both small molecules bind along the long axis of Aβ fibrils.

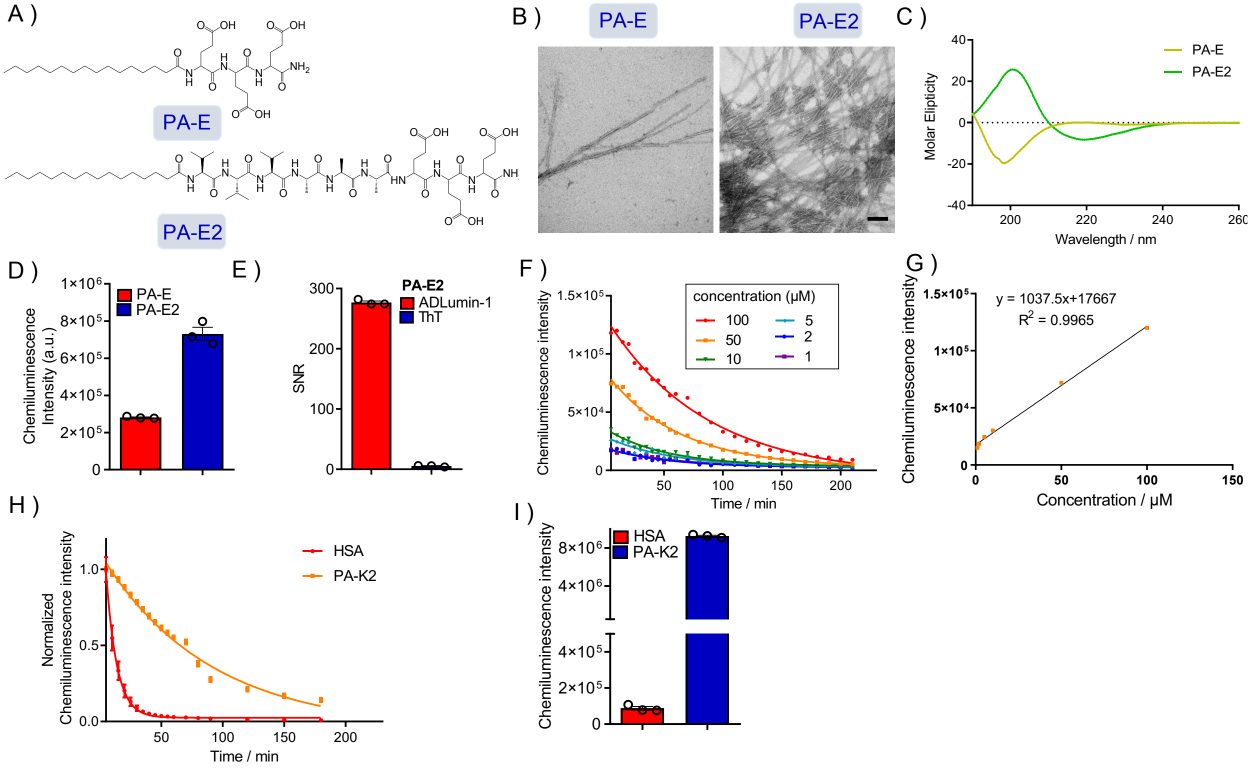


**Fig. S2.** ADLumin-1's specificity to β-sheet-rich structures. A) Chemical structure of PA-E/E2. B) Representative TEM images for PA-E/E2 (500 μM). C) CD spectra of PA-E/E2 (10 μM). PA-E2 showed typical peaks indicating the formation of β-sheet-rich structure while PA-E showed random coil structure. D) Chemiluminescence intensity of ADLumin-1 in the presence of PA-E/E2 (500 μM). E) SNR of ADLumin-1 and ThT with PA-E2 (500 μM). F) Chemiluminescence signal profile of ADLumin-1 with different concentrations of PA-E2 (1-100 μM) from 5 to 200 min. G) Linear regression of the highest signal of ADLumin-1 with different concentrations of PA-E2 (1-100 μM). H,I) Normalized signal decay profile of 500 μM human serum albumin (HSA) or PA-K2 after mixing with ADLumin-1 (H) and the quantified chemiluminescence intensity at 30 min (I). The slow decay in the presence of β-sheet-rich structure indicates that ADLumin-1 forms a strong and specific bond with the hydrophobic tunnel within the β-sheet-rich structure, whereas its interaction with albumin appears to be transient and non-specific.

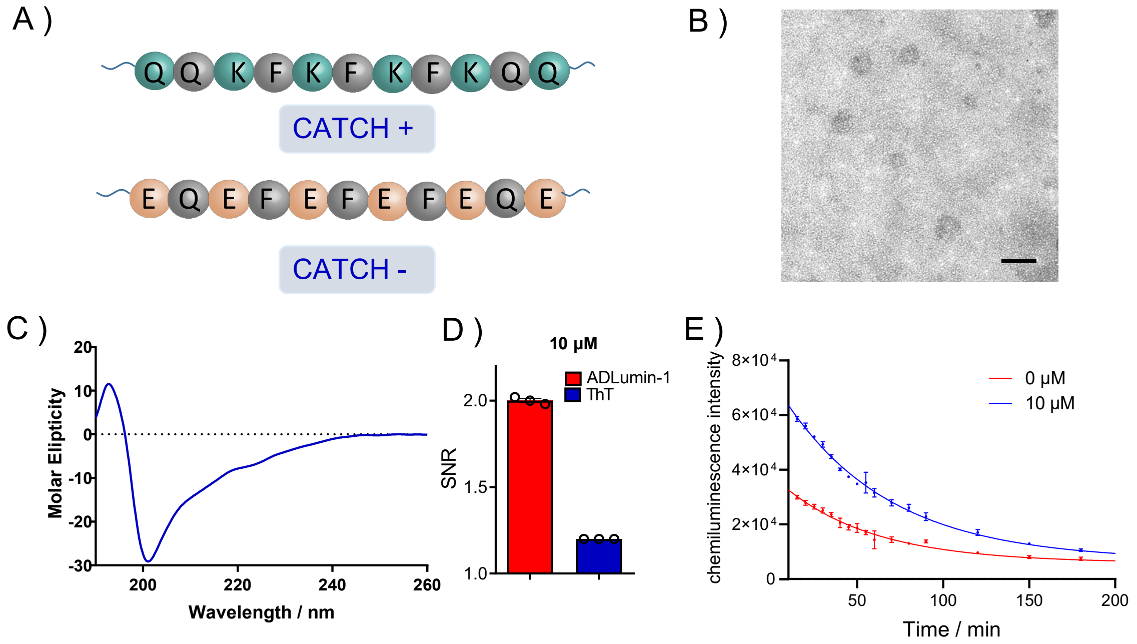


**Fig. S3.** Detecting capability of ADLumin-1 to oligomers. A) Chemical structure of CATCH+ and CATCH- peptide. B) Representative TEM images for CATCH+/- pairs (10 μM). Scale bar: 200 nm. C) CD spectra of CATCH+/- pairs. D) Signal-to-noise ratio (SNR) of ADLumin-1 in the presence of CATCH+/- pairs. E) Chemiluminescence signal profile of ADLumin-1 with or without CATCH+/- pairs (10 μM) from 5 to 200 min.

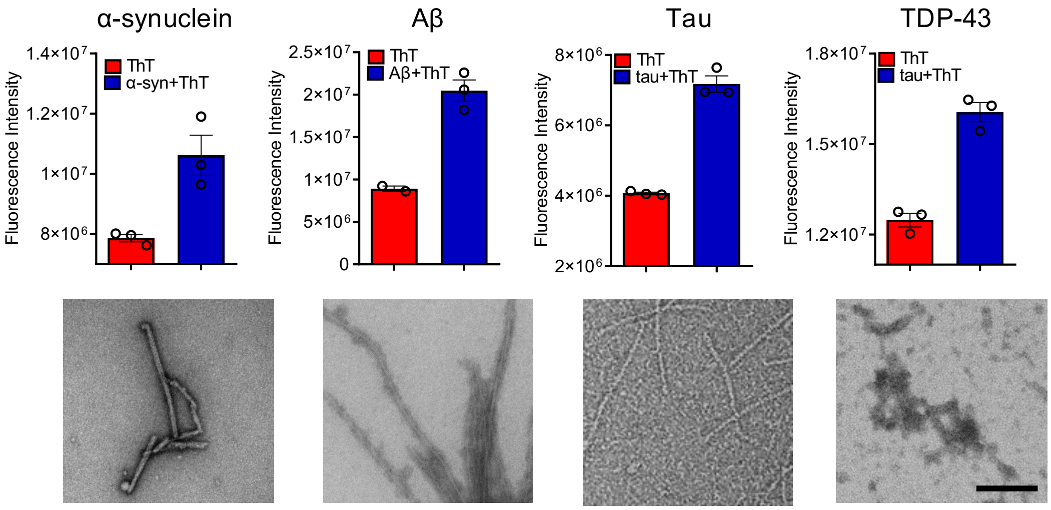


**Fig. S4.** Fibrils formation confirmation. The fibrils formation of misfolded proteins were confirmed by ThT fluorescence assay and TEM imaging. Scale bar: 200 nm.

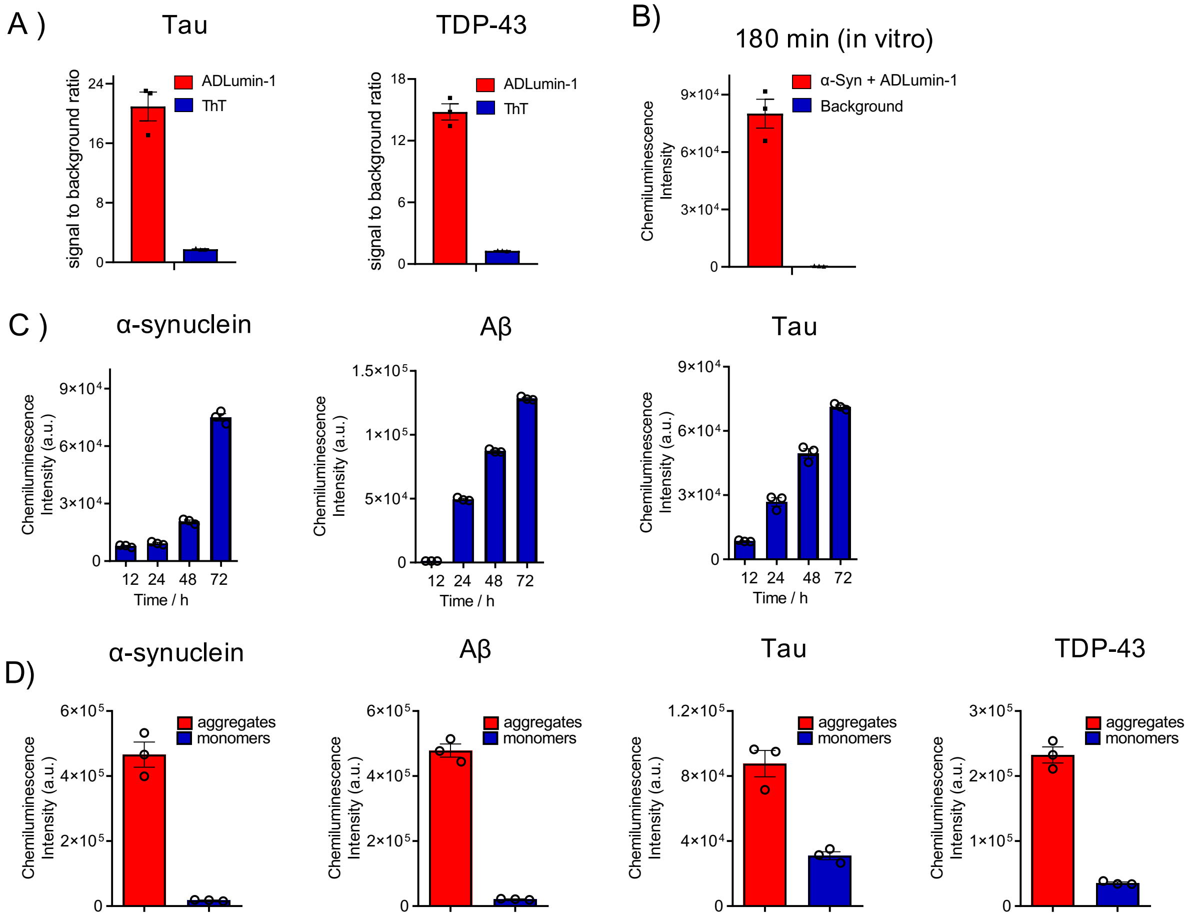


**Fig. S5.** ADLumin-1 could detect various misfoldons. A) Signal to background ratio of ADLumin-1 or ThT (final concentration: 15 µM in 5% DMSO/PBS buffer) when interacting with tau or TDP-43 aggregates. B) Chemiluminescence intensity of background or ADLumin-1 (final concentration: 15 µM in 5% DMSO/PBS buffer) when mixing with 70 µM α-syn aggregates and measured at 180 min. ADLumin-1 group showed an SNR of 222.15 ± 29.71 at 180 min, indicating long duration of chemiluminescence signal in vitro. C) Chemiluminescence intensity of ADLumin-1 for detecting α-syn, Aβ, tau, and TDP-43 after incubating and shaking for 12-72 h. With the increased degree of fibrils, the probe showed gradually increased signals. D) Chemiluminescence intensity of ADLumin-1 (final concentration: 15 µM in 5% DMSO/PBS buffer) when interacting with α-syn, Aβ, tau, and TDP-43 aggregates or monomers. ADLumin-1 showed signal increase for aggregated proteins but weak signals for monomeric forms.

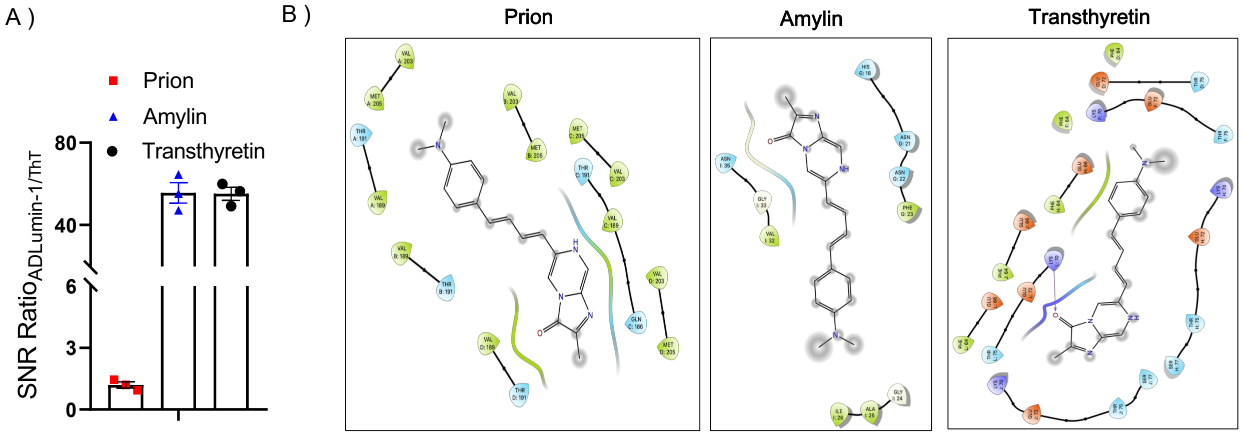


**Fig. S6.** The interaction of ADLumin-1 to prion, amylin and transthyretin. A) The SNR ratio of ADLumin-1 to ThT (final concentration: 15 μM in 5% DMSO/PBS buffer) when mixing with misfolded prion, amylin and transthyretin. ADLumin-1 showed significantly higher signal SNR compared to ThT when interacting with misfoldons. B) Key interacting residues of ADLumin-1 to misfoldons. ADLumin-1 interacts with the hydrophobic tunnels that are formed by Ala, Val, Ile or Phe tested by molecular docking.

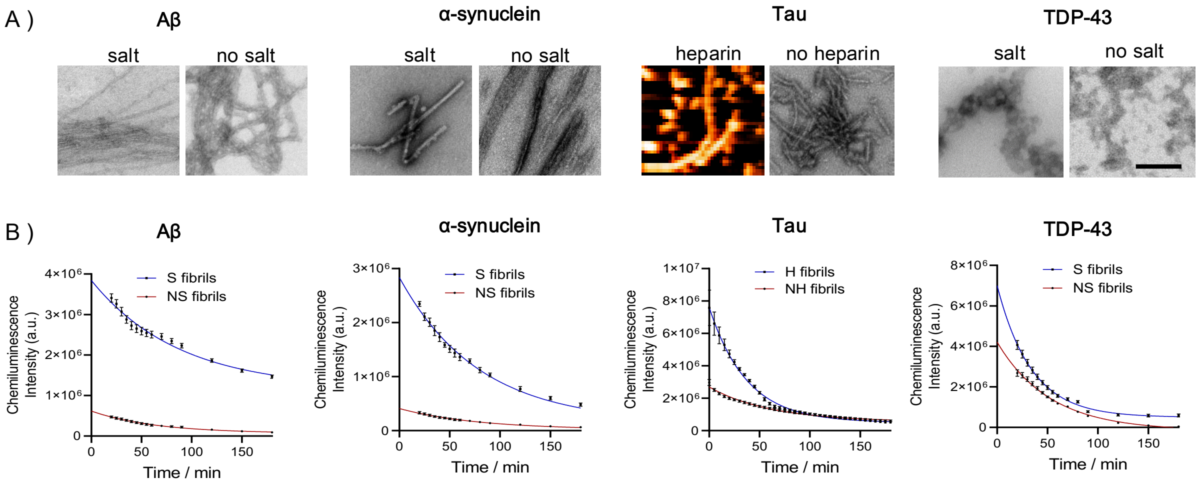


**Fig. S7.** The interaction of ADLumin-1 to fibril polymorphs. A) TEM/AFM imaging of misfolding-prone proteins that fibrillated under salt (sodium chloride, S; heparin, H) or non-salt (no sodium chloride, NS; no heparin, NH) conditions. The proteins displayed distinct morphologies with or without salt/heparin. Scale bar: 200 nm. B) The chemiluminescence signal of 15 μM ADLumin-1 after mixing with different fibrils from 0 to 180 min. ADLumin-1 showed distinct intensity and decay profiles for different fibrils.

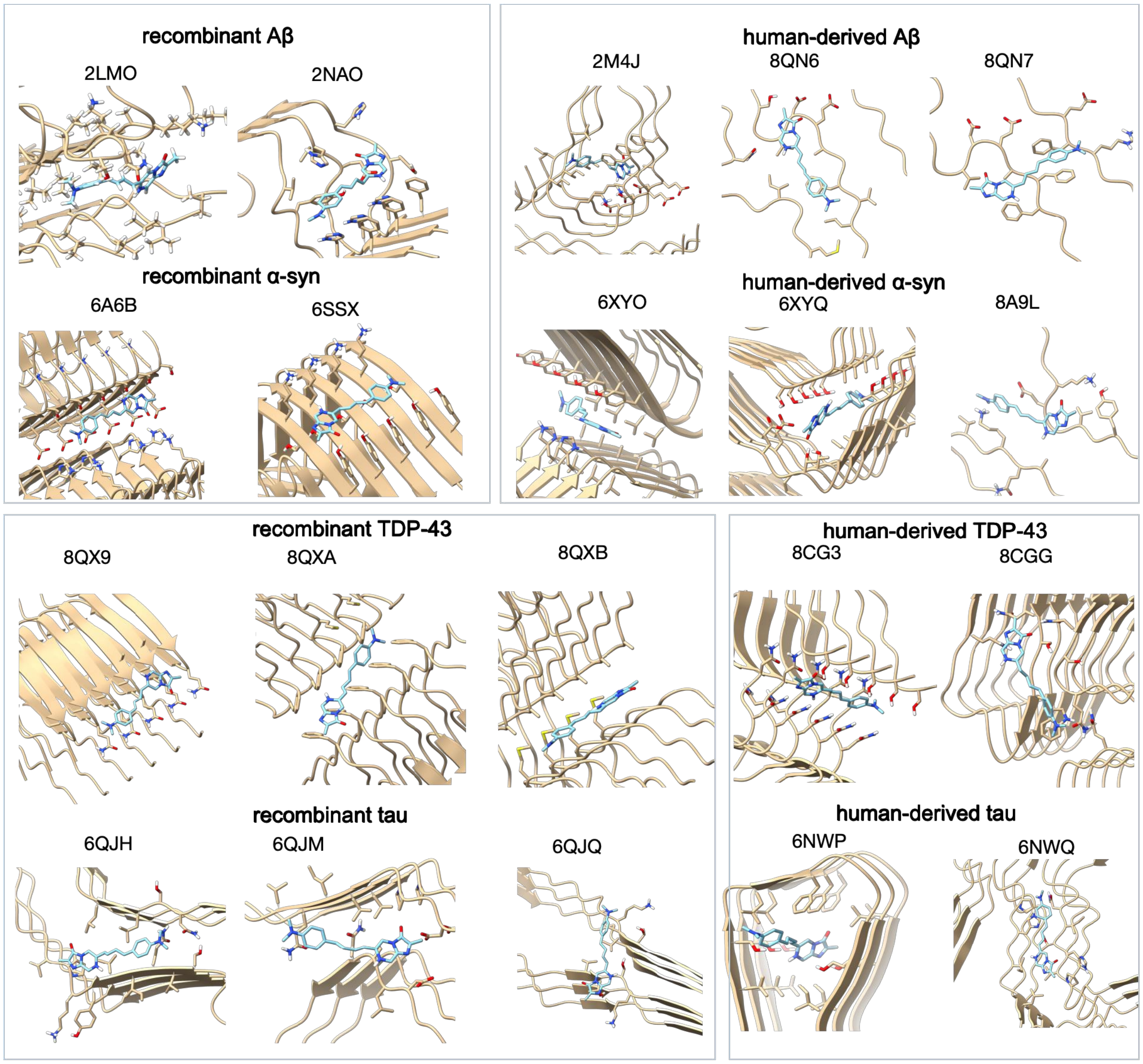


**Fig. S8.** Molecular docking of ADLumin-1 with various recombinant and human brain-derived fibrils.

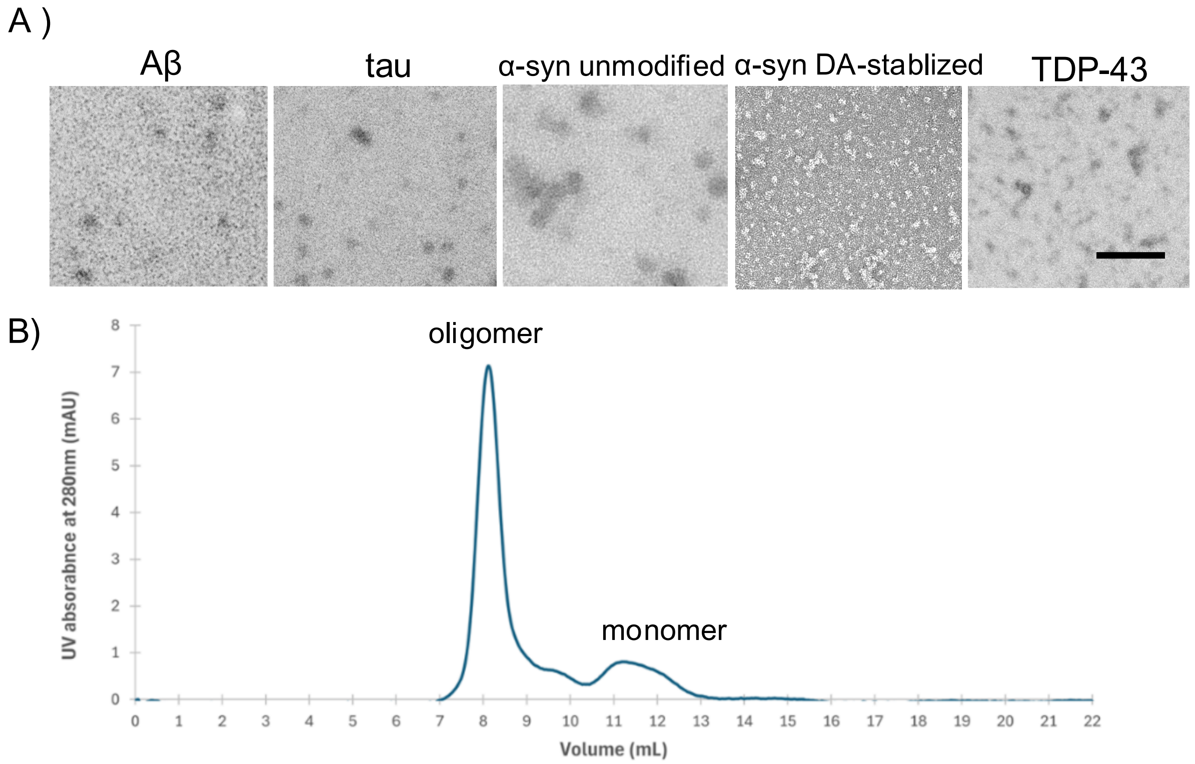


**Fig. S9.** A) Representative TEM images of misfolded protein oligomers. Scale bar: 200 nm. B) Representative SEC images of oligomer purification.

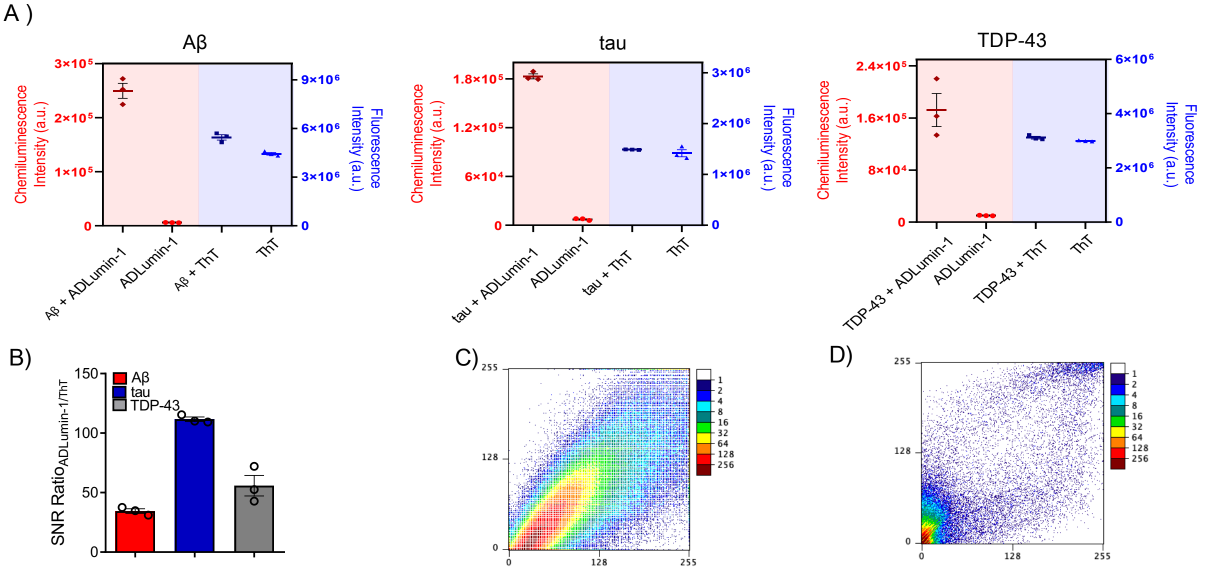


**Fig. S10.** A, B) The chemiluminescence or fluorescence signal of 15 μM ADLumin-1 or ThT when mixing with misfolded protein oligomers. ADLumin-1 showed significantly higher signal enhancements (A) and SNR (B) compared to ThT. C, D) Colocalization analysis of ADLumin-1 with anti-tau antibody staining in Alzheimer's disease patient brain tissue slides (C), and with anti-TDP-43 antibody staining in A315T transgenic mice tissue slides (D). ADLumin-1 showed good colocalization with both antibody stainings and Pearson's R value was 0.88 and 0.81, respectively.

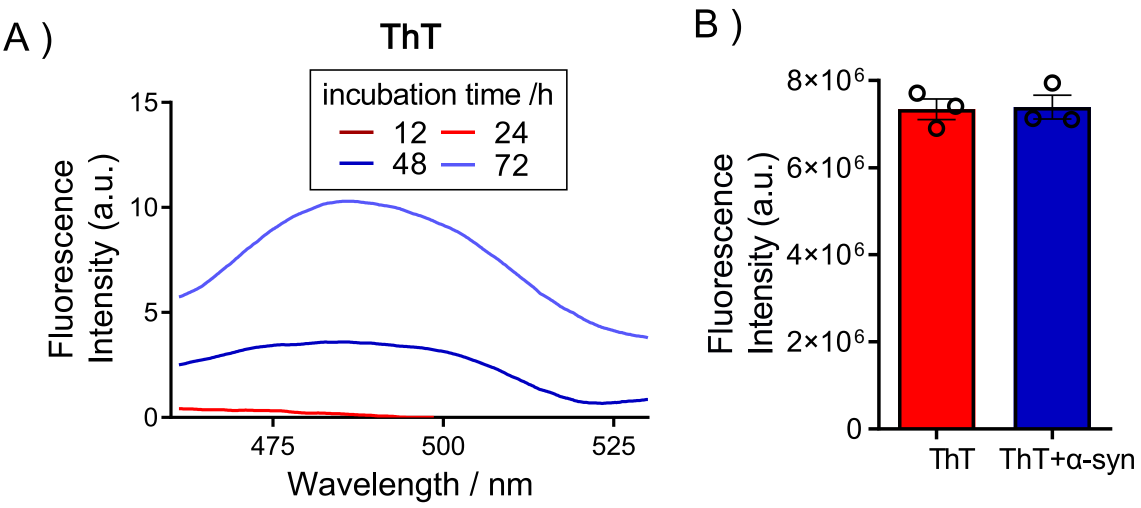


**Fig. S11.** Detecting capability of ThT to α-syn. A) Monitoring α-syn aggregation (2.5 μM) with increased incubation time (12-72 h) by using ThT. B) ThT fails to directly detect 1 pg/mL α-syn fibrils in PBS buffer.

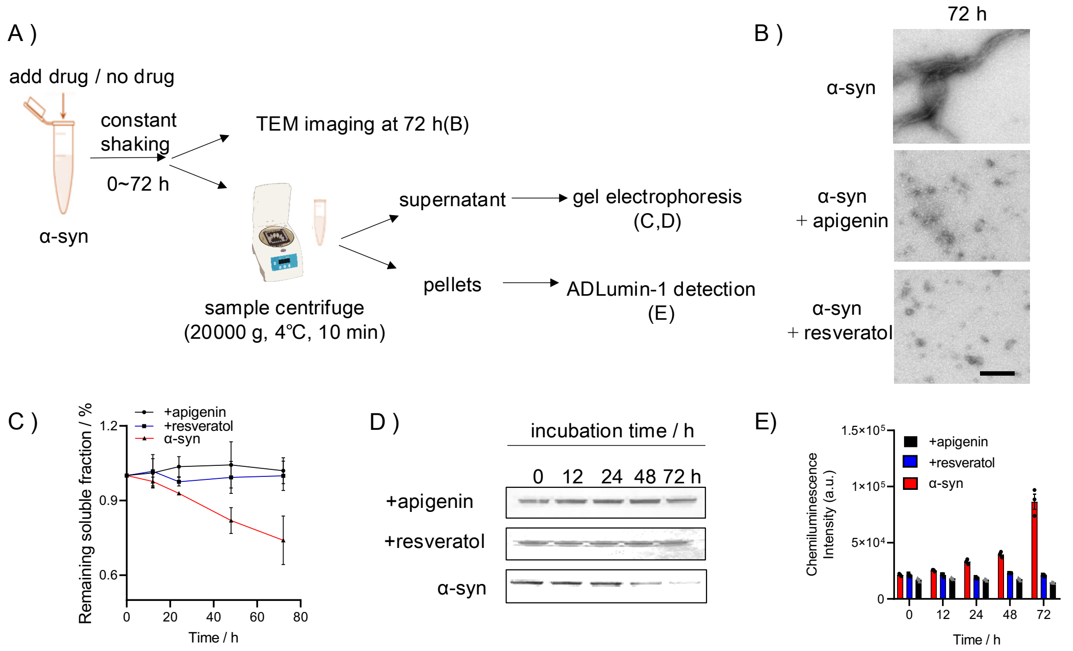


**Fig. S12.** A) Diagram of the workflow of sedimentation assay. B) Representative TEM images of α-syn samples with or without inhibitor treatment and incubated under shaking for 72 h. We found fibrils in α-syn group but soluble species in the inhibitor-treated group. Scale bar: 200 nm. C) Remaining soluble fraction (%) quantified by grey intensity of gel electrophoresis results. The drug treated group showed negligible change in soluble species while α-syn only group showed drastically decreased soluble species. D) Representative gel electrophoresis images of α-syn with or without drug treatment from 0 to 72 h. E) Chemiluminescence intensity of ADLumin-1 with insoluble α-syn samples in sedimentation experiments. α-syn proteins showed increased signal during 12-72 h. With the inhibitors (Apigenin or Resveratrol), the signal showed negligible changes over time, indicating the inhibitors significantly inhibited the formation of α-syn fibrils, and this process can be detected by ADLumin-1.

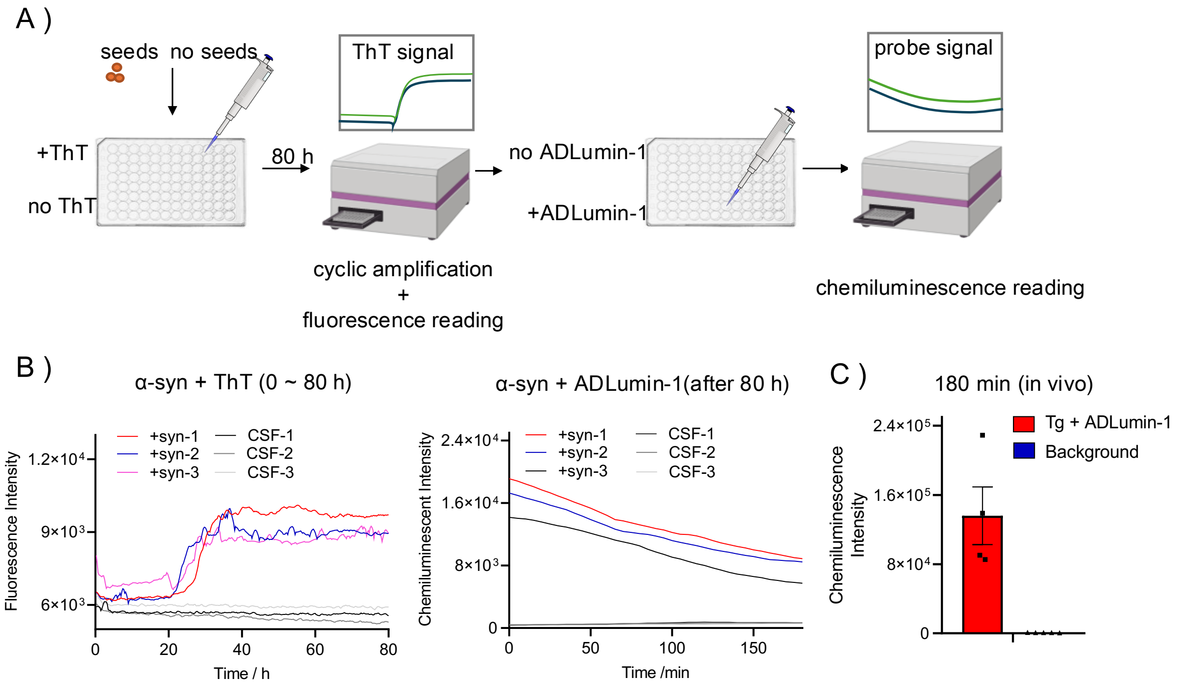


**Fig. S13.** A) Diagram of the workflow of protein cyclic amplification by using a microplate reader. As chemiluminescence is different from fluorescence, the form is not reversible upon emitting light, while fluorescence is reversible and can be repeatedly excited. ThT was added before the experiments and monitored from 0 to 80 h under cyclic shaking and incubation, and ADLumin-1 was added after the cyclic amplification. B) Fluorescence intensity of ThT or chemiluminescence signal decay profile of ADLumin-1 when incubated with CSF samples with or without 35 pM α-syn pre-fibrillated seeds in the cyclic amplification experiments. By comparing the highest intensity, we found ADLumin-1 showed 32.56 ± 5.79-fold higher signal compared to ThT. C) Chemiluminescence intensity of background or ADLumin-1 when i.p. injected into transgenic A53T mice and measured at 180 min postinjection. ADLumin-1 group showed SNR of 177.30 ± 75.38 at 180 min, indicating long duration of chemiluminescence signal in vivo.

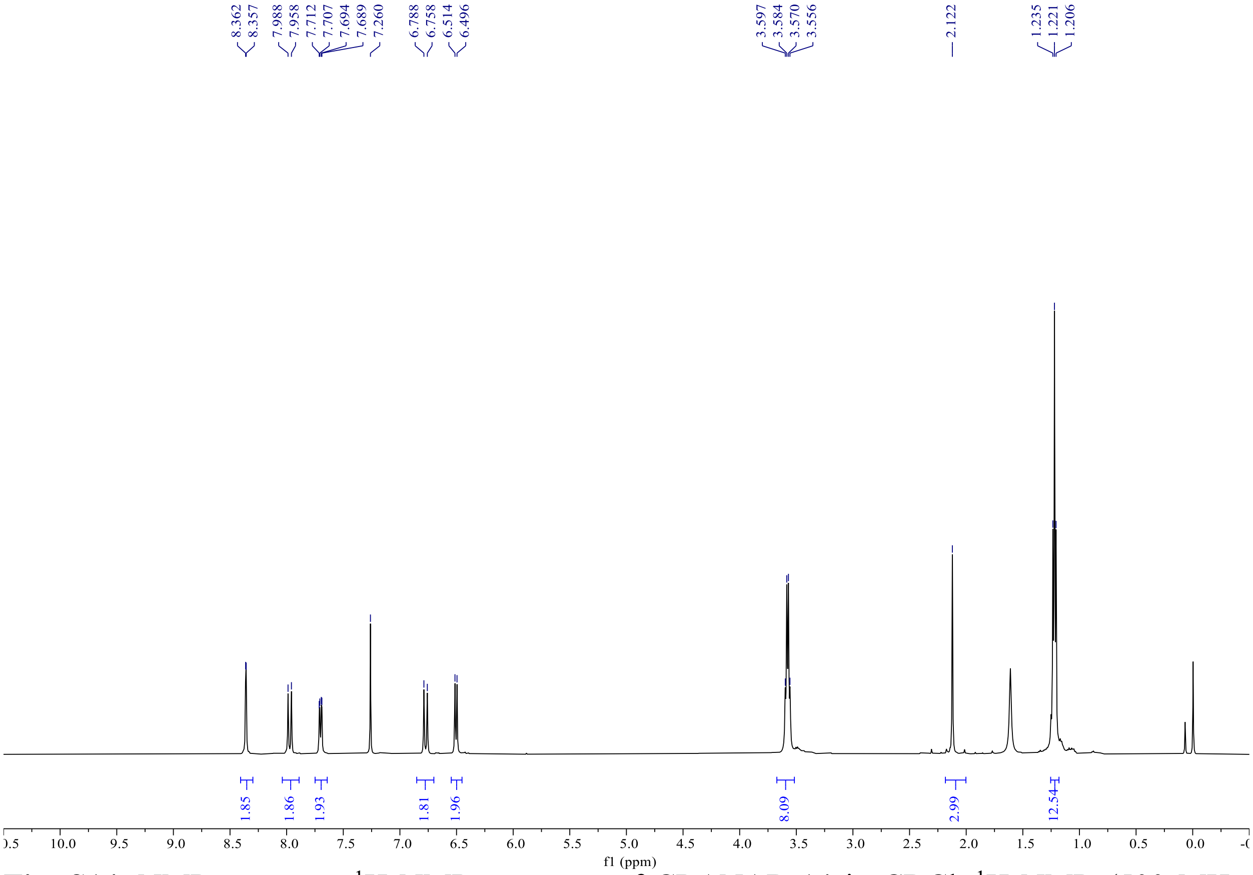

**Fig. S14.** NMR spectrum $^1H$ NMR spectrum of CRANAD-14 in $CDCl_3$.$^1H$ NMR (500 MHz, $CDCl_3$) δ 8.36 (d, J = 2.4 Hz, 1H), 7.97 (d, J = 15.0 Hz, 1H), 7.70 (dd, J = 9.1, 2.5 Hz, 1H), 6.77 (d, J = 15.1 Hz, 1H), 6.50 (d, J = 9.0 Hz, 1H), 3.58 (q, J = 6.7 Hz, 5H), 2.12 (s, 2H), 1.22 (t, J = 7.1 Hz, 7H).

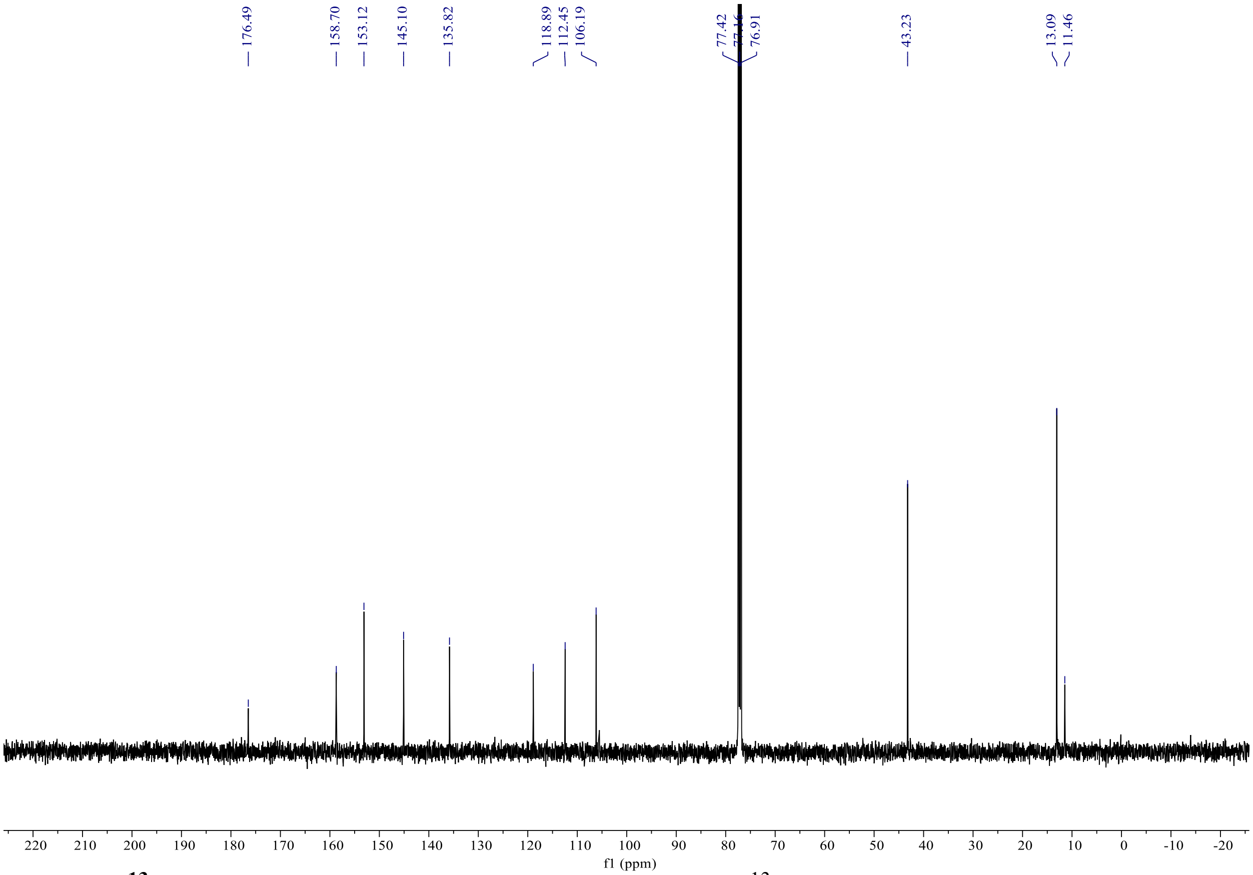


**Fig. S15. $^{13}$C NMR spectrum of CRANAD-14 in $CDCl_3$.** $^{13}$C NMR (126 MHz, CDCl3) δ 176.49, 158.70, 153.12, 145.10, 135.82, 118.89, 112.45, 106.19, 43.23, 13.09, 11.46.

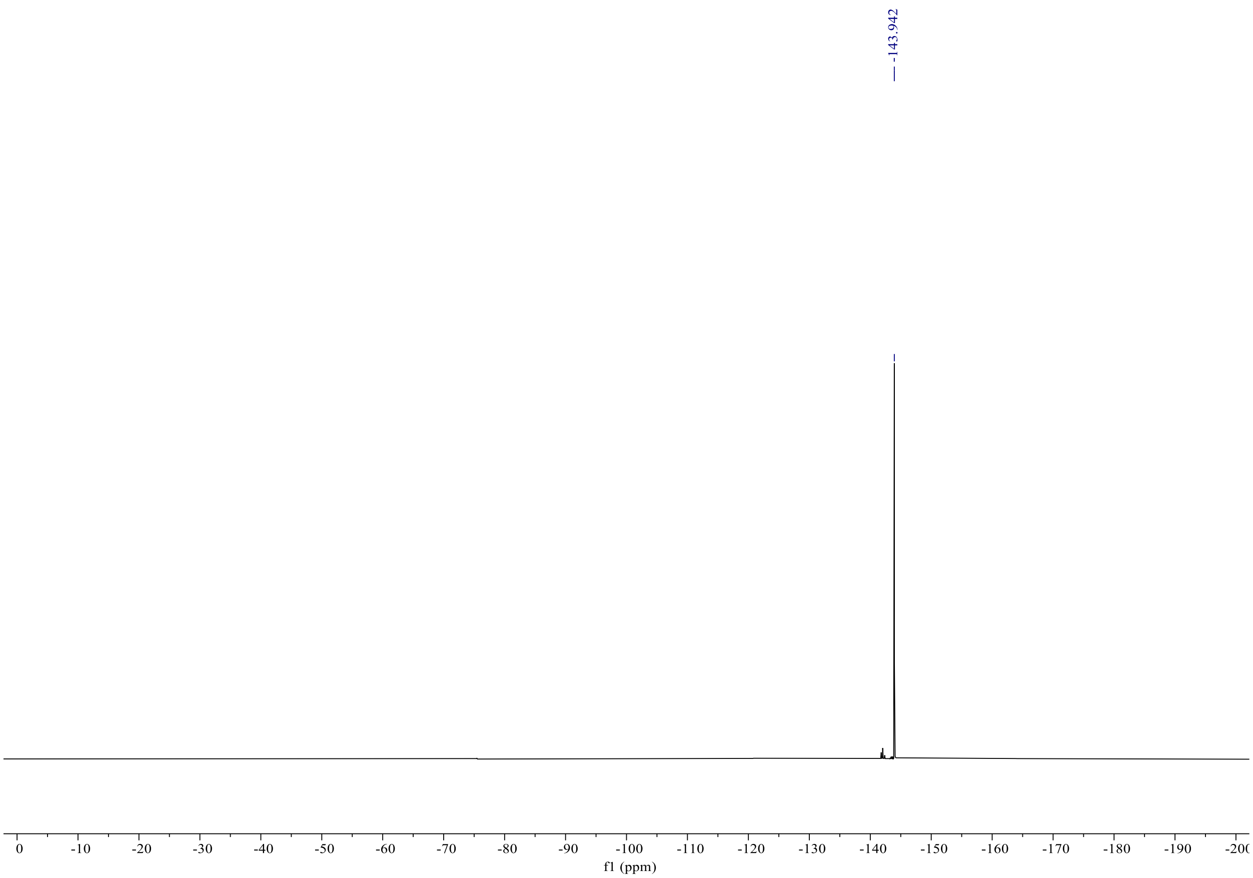


**Fig. S16. $^{19}$F NMR spectrum of CRANAD-14 in $CDCl_3$.** $^{19}$F NMR (470 MHz, CHLOROFORM-D) δ -143.94.

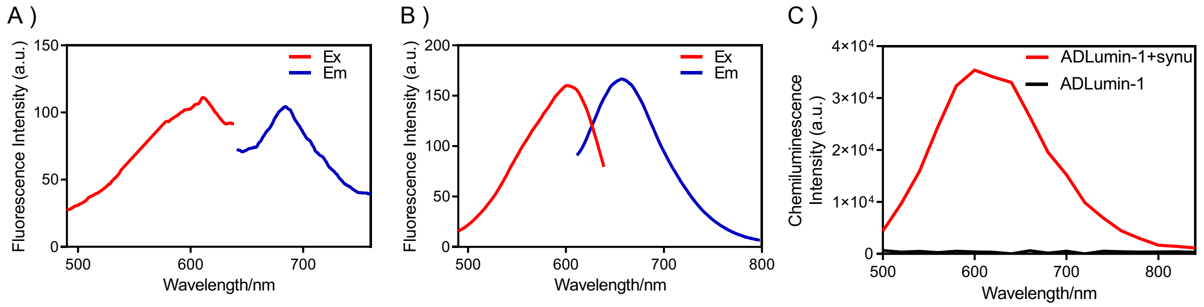


**Fig. S17.** Optical spectra of CRANAD-14 and ADLumin-1. A, B) Fluorescence spectra of CRANAD-14 before and after interacting with the α-synuclein aggregates. CRANAD-14 alone showed excitation wavelength at 611 nm, and emission wavelength at 685 nm (A). After interacting with α-synuclein aggregates, the fluorophore showed excitation wavelength at 604 nm, and emission wavelength at 661 nm (B). C) Chemiluminescence spectra of ADLumin-1 with or without α-synuclein aggregates. The results showed good overlap between the chemiluminescence spectra of ADLumin-1 and excitation spectra of CRANAD-14 and thus enable ChRET imaging.

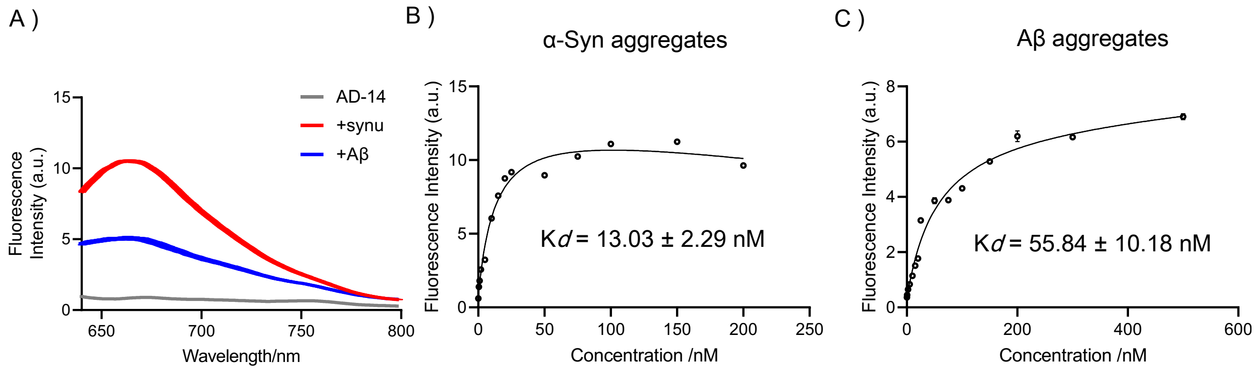


**Fig. S18.** Selectivity of CRANAD-14 for α-synuclein and Aβ aggregates. Fluorescence intensity (A) and dissociate constant of CRANAD-14 with 5 μM α-synuclein aggregates (B) or Aβ aggregates (C).

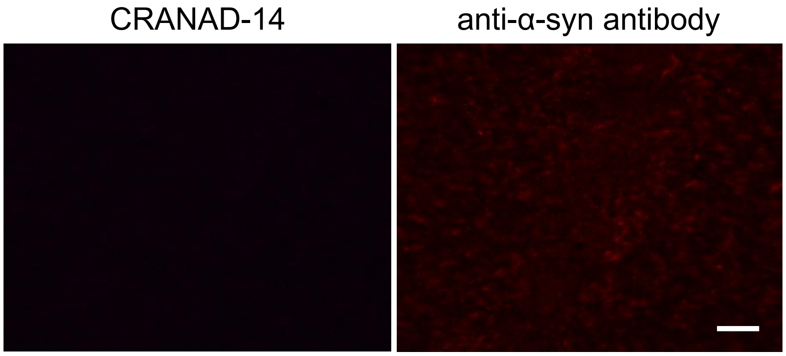


**Fig. S19.** Ex vivo histological observation of wild-type mice (female, 9-month-old, n = 3) after CRANAD-14 injection. The brain tissue slides of wild-type mice showed no obvious labeling (scale bar = 20 μm).

**Table S1.** Signal decay half-life time of ADLumin-1 with various misfoldons.

| Protein (PDB: ID) | Half-life Time / min |
|---|---|
| Prion | 29.21 |
| Amylin | 23.22 |
| Tau | 22.34 |
| Alpha-Synuclein | 93.03 |
| Amyloid Beta | 49.04 |
| Transthyretin | 22.21 |
| TDP-43 | 33.42 |

**Table S2.** Docking scores and interacting moieties of ADLumin-1 to various misfoldons.

| Protein (PDB: ID) | Best Score (kcal/mol) | AA Segment (Chain: AA) |
|---|---|---|
| Prion (6LNI) | -9.7 | A-J: KE |
| Amylin (6VW2) | -6.4 | A,D,E,I: SST; H: S |
| Tau (5O3L) | -7.3 | A,E,G: RVQ; C,I: RQ |
| Alpha-Synuclein (6PEO) | -8.8 | A,C: AKTV; B,E: TAKTV; D: TV |
| Amyloid Beta (5OQV) | -9.7 | A: FNKI; C,H: FNKAI; E: KAI; F: F NAI |
| Transthyretin (7OB4) | -7.2 | D,F: EFK; H: F; J: FKETS; L: ETS |
| TDP-43 (7PY2) | -7.1 | A,D: GFGNQ; B,C: GFGNQG |

**Table S3.** Docking scores and interacting moieties of ADLumin-1 to fibrils polymorphs.

| | | PDB ID | Score (kcal/mol) |
|---|---|---|---|
| Aβ | recombinant | 5OQV | -9.7 |
| | | 2LMO | -7.606 |
| | | 2NAO | -5.78 |
| | human-derived | 2M4J | -6.836 |
| | | 8QN6 | -5.398 |
| | | 8QN7 | -5.66 |
| α-syn | recombinant | 6PEO | -8.8 |
| | | 6A6B | -8.417 |
| | | 6SSX | -6.877 |
| | human-derived | 6XYO | -9.123 |
| | | 6XYQ | -8.787 |
| | | 8A9L | -4.966 |
| tau | recombinant | 6QJH | -5.273 |
| | | 6QJM | -5.905 |
| | | 6QJQ | -3.56 |
| | human-derived | 5O3L | -7.3 |
| | | 6NWP | -6.842 |
| | | 6NWQ | -3.979 |
| TDP-43 | recombinant | 8QX9 | -6.779 |
| | | 8QXA | -6.959 |
| | | 8QXB | -7.346 |
| | human-derived | 7PY2 | -7.1 |
| | | 8CG3 | -8.546 |
| | | 8CGG | -5.76 |
